# A mushroom-body output neuron that mediates octopamine-driven and hunger-motivated feeding in *Drosophila*

**DOI:** 10.64898/2026.03.13.711740

**Authors:** Xianyuan Zhang, Sangyu Xu, Joses Ho, James C. Stewart, Adam Claridge-Chang

## Abstract

Feeding behavior requires the integration of environmental cues, metabolic signals, and internal states through neuromodulatory networks, yet the role of specific neuromodulatory neurons and how they are coordinated in controlling feeding remains unclear. Using automated feed-tracking and optogenetics in *Drosophila melanogaster*, we found that activation of two octopaminergic neurons (VPM3/4) increases food consumption. Both VPM neurons form direct synapses with a mushroom-body output neuron MBON11, which, through opto-activation and opto-inhibition, can exert bidirectional control over food intake, with sexually dimorphic effects. Epistasis experiments demonstrated that octopamine-driven feeding requires functional MBON11 output. Additionally, we found that the dopaminergic PPL101 neurons, which also synapse with MBON11, are required for hunger-driven feeding. Ethomic analysis contextualized with natural hunger and satiety provided a holistic view of how different neuron types linked to the mushroom-body influence feeding behaviors. This analysis revealed that MBON11 interventions best recapitulate natural hunger-satiety transitions. These findings revealed a circuit where two neuromodulatory neuron types with distinct unidirectional feeding effects—octopaminergic VPM3/4 (instructive but not required) and dopaminergic PPL101 (required but not instructive)—converge onto MBON11, a neuron whose activity is both required for and instructive of hunger-related feeding. This circuit arrangement may represent an architecture for integrating multiple motivational signals in feeding regulation.

**Key points:**

1. When activated, octopaminergic VPM3 and VPM4 cells drive increased food consumption, effects that require octopamine signaling and are dependent on MBON11 function.
2. Octopaminergic neuronal activity is not required for hunger-driven feeding.
3. Receiving direct synaptic inputs from VPM3 and 4, MBON11 neurons bidirectionally control feeding, and broadly phenocopy natural hunger–satiety transitions.
4. Dopaminergic PPL101 neurons that synapse onto MBON11 are required for hunger-motivated feeding.
5. MBON11 appears to integrate octopaminergic and dopaminergic signals, and regulate the feeding behavioral state.

## Introduction

Feeding is fundamental to animal survival, with evidence suggesting that several hunger and satiety genes, the sensory-motor system and motivational systems that regulate feeding behavior are conserved across species (Tinbergen 1989; Tierney 1995; Yarmolinsky et al. 2009; Lin et al. 2019; Miroschnikow et al. 2020; Betley et al. 2013). Dysfunctions in these systems can lead to psychiatric, neurological, and metabolic conditions (Morganstern et al. 2011; Hoek 2013; Lee and Blackshaw 2014; Naef et al. 2015), including obesity, which has emerged as a major global-health burden over the past decade (GBD 2021 Diseases and Injuries Collaborators 2024). Feeding behavior comprises a sequence of actions, including nutrient interoception, food detection and location, feeding initiation and termination, and digestion (Pool and Scott 2014; Kim et al. 2017; Wang and Wang 2019). These actions are influenced by external stimuli, such as food odors, and the internal states, notably hunger and satiety (Watts et al. 2022). The vinegar fly, *Drosophila melanogaster*, offers a genetically tractable model to dissect the molecular and neural mechanisms underlying these processes (Itskov and Ribeiro 2013).

Nutritional states are intricately regulated by evolutionarily ancient chemical messengers known as neuromodulators (Moroz et al. 2021; Goulty et al. 2023; Bargmann 2012). In mammals, adrenergic signaling in the brain, mediated by norepinephrine-releasing cells, has been investigated for its role in regulating feeding (Leibowitz 1975; Rossi et al. 1982; Adan et al. 2008; Hostrup and Onslev 2022; Sayar-Atasoy et al. 2023), however, the specific contributions of individual central adrenergic neurons remain poorly understood. Invertebrates, including flies, lack norepinephrinergic circuits and instead use octopamine (OA), a neuromodulator with similar physiological functions (Roeder 2005). In flies, OA has been found to mediate hunger-induced hyperactivity associated with food-seeking behavior (Yang et al. 2015). OA was also found to modulate sugar responsiveness (Scheiner et al. 2014; Damrau et al. 2017), signal sweetness, and induce short-term sugar-conditioned appetitive memory (Burke et al. 2012; Huetteroth et al. 2015). Through interactions with insulin signaling, octopamine has also been implicated in metabolic regulation (Yu et al. 2016; Held et al. 2025; Berger et al. 2024; Li et al. 2016; Bisen et al. 2025). The diverse (and sometimes opposing) roles of a neuromodulator can often be attributed to its release by distinct cell types (Chandler et al. 2014; Uematsu et al. 2015; Claßen and Scholz 2018). Understanding how OA regulates feeding therefore requires examining specific OA neuronal sets.

Specific ventromedial octopaminergic neurons located in the subesophageal zone of the fly brain have been associated with sugar reward and feeding-related behaviors (Hammer 1993; Burke et al. 2012; Yu et al. 2016; Hermanns et al. 2022). Among these neurons, the ventral-paired-medial 4 (VPM4) class has been implicated in feeding-associated behaviors. VPM4 neurons have been found to signal nutritional needs and promote proboscis extension (Youn et al. 2018). Another study associated VPM4 neurons with food reward and the inhibition of persistent food-odor tracking (Sayin et al. 2019). Summarizing these observations, a review described the function of VPM4 as the neuronal directive “stop looking, start eating” (Selcho and Pauls 2019). Previous studies have examined feeding-associated proxies, so the direct role of VPM4 on regulating food consumption *per se* remains unaddressed. Like VPM4, the VPM3 major neurites first arborize in the esophagus, traverse the posterior margin of the antennal lobes, and project axons to various regions in the superior protocerebrum, including the γ lobe and peduncle of the mushroom body (MB) (Busch et al. 2009; Aso, Hattori, et al. 2014). Recent studies have demonstrated that VPM3 plays a role in prioritizing recent food reward memories over remote ones (Kapoor and Waddell 2024), and also regulates sleep and courtship in a sex-dependent manner (Reyes et al. 2025).

The projection of VPM neurons to the MB suggests a potential circuit mechanism for feeding regulation. The MB has long been recognized as the primary learning and memory center in insects (Heisenberg et al. 1985; McGuire et al. 2001; Dubnau et al. 2001). However, recent studies have expanded this view, revealing that the MB also functions as a neural integration hub that combines diverse sensory inputs with internal state information to modulate both sensory and motivation-driven behavioral decisions such as food seeking (Tsao et al. 2018a; Suárez-Grimalt et al. 2024). Recent connectomics studies have revealed large synaptic outputs from both VPM3 (80 connections) and VPM4 (83 connections) neurons to mushroom body output neuron 11 (MBON11), also referred to as the MBON-γ1pedc>α/β cells and the MB-MVP2 cells (Tanaka et al. 2008; Scheffer et al. 2020). The VPM4→MBON11 connection has attracted interest, with studies showing that MBON11 has roles in regulating feeding-related innate and learned olfactory behavior (Aso, Sitaraman, et al. 2014; Perisse et al. 2016; Tsao et al. 2018a; Sayin et al. 2019; Matheson et al. 2022). MBON11 cells are GABAergic and project axons across the MB’s entire α/β lobes, the γ4 and γ5 zones: regions associated with short-term olfactory and visual memories, acute valence, naive odor avoidance/approach, and olfactory navigation (Li et al. 2020; Aso, Sitaraman, et al. 2014; Owald et al. 2015; Matheson et al. 2022). However, these previous studies have not directly measured food consumption or tested the role of MBON11 activity in controlling feeding behavior *per se*.

The MBON11 neurons receive substantial input (MBON11 in the right hemisphere receives 1,950 connections in total from both hemispheres) from another set of neuromodulatory cells: MB-projecting dopaminergic paired-posterior lateral neurons PPL101 (also known as the PPL1-γ1pedc or MB-MP1 cells, based on their MB zonal projections) (Scheffer et al. 2020). These neurons have been found to signal punishment in olfactory, taste, and visual learning (Schwaerzel et al. 2003; Keene and Waddell 2007; Claridge-Chang et al. 2009; Mao and Davis 2009; Vogt et al. 2014; Masek et al. 2015). Several studies have suggested that PPL101 neurons signal satiety (Krashes et al. 2009; Meschi et al. 2024) and that diminished PPL101 activity promotes food seeking (Tsao et al. 2018a), whereas another study demonstrated that inhibiting PPL101 impairs food-odor tracking (Sayin et al. 2019). While PPL101 neurons have been implicated in feeding-related behaviors, whether PPL101 signals hunger, satiety, and consumption is also unclear.

Many *Drosophila* feeding-related assays examine behavioral sub-programs such as proboscis extension, odor tracking, and food contact (Shiraiwa and Carlson 2007; Itskov et al. 2014; Ro et al. 2014; Tsao et al. 2018a; Sayin et al. 2019), or measure total consumption over time without real-time tracking, such as the CApillary FEeder Assay (CAFE) (Ja et al. 2007; Deshpande et al. 2014). Recent automated capillary systems include Expresso (Yapici et al. 2016; Whitehead et al. 2024) and the Activity-Recording CAFE (ARC) (Murphy et al. 2017). We developed a video-tracked capillary-feeding apparatus, termed “Espresso”, that can be used to simultaneously monitor food consumption, track locomotion, and also facilitate optogenetic interventions. This apparatus thus provides multi-dimensional behavioral data suitable for ethomic analysis.

In this study, we aimed to investigate the roles of the MB-projecting OA neurons VPM3 and VPM4, as well as MBON11 and PPL101 cells, in regulating *Drosophila* food consumption. We used the Espresso apparatus to examine various behavioral features of feeding *per se*. By combining behavioral assays, optogenetic actuation, and ethomic analysis, we sought to provide a holistic view of how interventions in four types of mushroom-body-projecting neurons influence feeding behaviors in the context of natural hunger and satiety.

## Results

An automated optogenetic feeding assay shows that Tdc2+ activity drives consumption We first hypothesized that OA neuron activation can induce hunger and promote feeding behavior, including food consumption. Using the Espresso assay (**Figure 1A-C**), we studied feeding behavior of flies expressing Chrimson (Chr), a red-light-sensitive channelrhodopsin (Klapoetke et al. 2014), under the control of the broad octopaminergic driver *Tdc2-Gal4*. Optogenetic activation of Tdc2 neurons during a 2-h experiment caused previously *ad libitum-*fed flies to feed more frequently and consume more food (**Figure 1D, E**). Additionally, the flies exhibited faster locomotion (**Figure 1F**). Detailed analysis of multiple feeding-behavior features revealed that Tdc2-cell activation induced an increase in feed volume, feed count, feed duration, meal size, and average locomotion speed, while decreasing latency before the first feed, thereby accelerating feeding initiation (**Figure 1G-H, Figure S1A-D**). In a similar experiment where food was omitted from the capillaries, flies showed a similar increase in locomotor speed, indicating that the observed optogenetically induced speed increase was not conditional on food odor or energy influx from nutrients (**Figure S1E-F**). These observations show that broad activation of all octopaminergic neurons promotes multiple foraging-like behaviors, including locomotion, feeding initiation, and food consumption.

**Figure 1.**
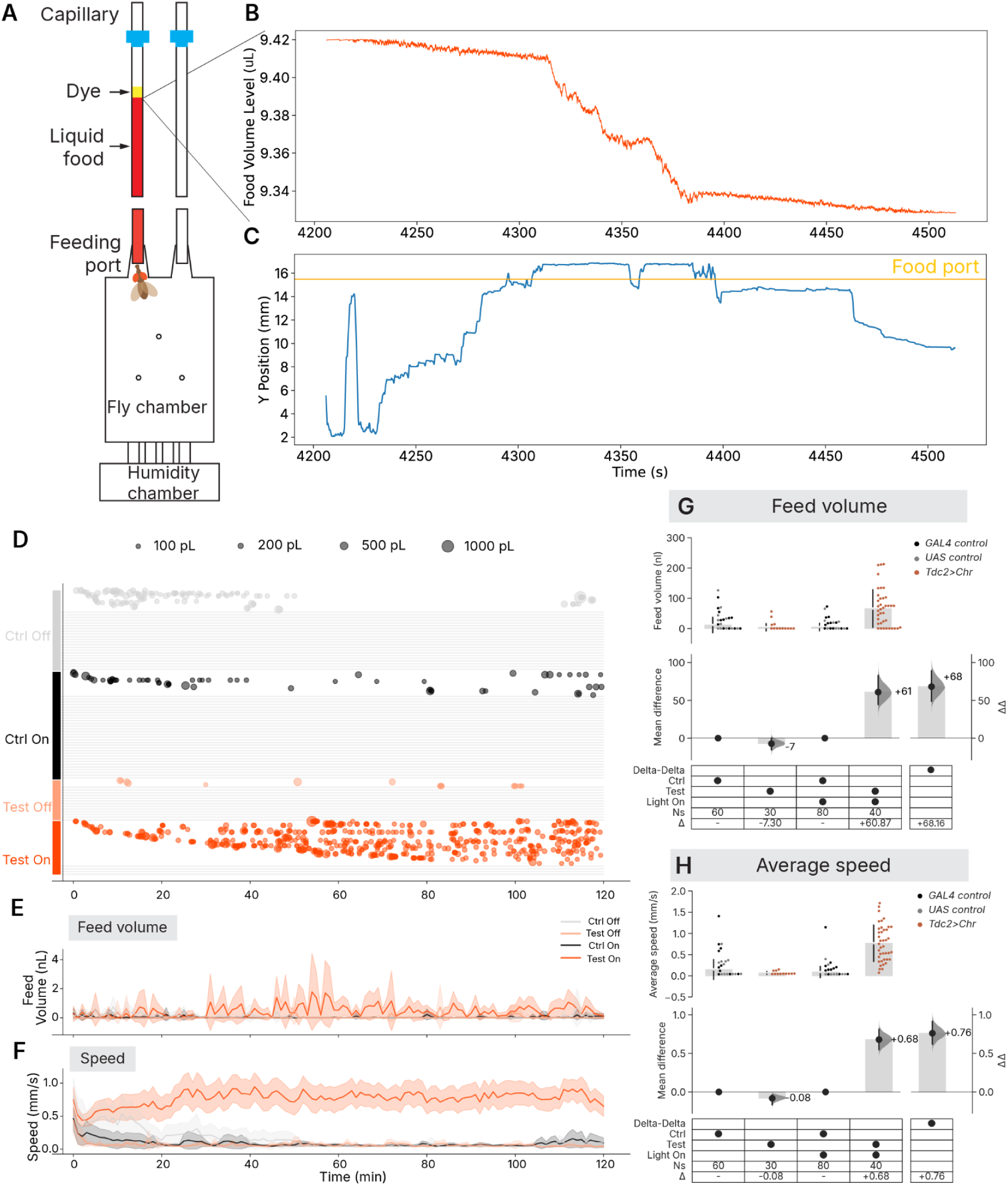
Automated capillary feeding assay reveals optogenetic activation of Tdc2+ neurons modulates feeding and locomotion. **A.** Diagram of a single Espresso unit, consisting of a capillary filled with liquid food (5 % w/v sucrose and 5 % w/v yeast extract, dyed for visualization), a feeding port, a fly chamber, and a humidity chamber. **B.** Representative trace of liquid-food level-tracked volume changes in a capillary during feeding events by a *Tdc2>Chr* fly. The food level in each capillary was continuously imaged and quantified throughout the assay. The rapid decline in food volume reflects food consumption. For more information see Methods. **C.** The Y-position (mm) of a fly during feeding events, corresponding to the same *Tdc2>Chr* fly and time window as in panel B. The fly’s vertical movement is tracked, with sustained periods at the food port relating to feeding. **D.** Bubble plot showing each feeding event (x-axis: time in minutes; y-axis: individual flies) and meal size for previously fed male *Tdc2>Chr* flies and genetic controls (driver and responder lines). Bubble area is scaled to volume, see key bubbles for 100 pL, 200 pL, 500 pL, 1000 pL. Rows represent individual flies, with “Ctrl On/Off” and “Test On/Off” indicating baseline/light-off conditions or red light activation (optogenetic stimulation). Non-feeding flies are indicated with alternating white/grey strips. **E.** Time series of average feed volume (nL) for fed male *Tdc2>Chr* flies and controls during a 2-h assay, under red light-on and light-off conditions. Lines represent group means, with shaded ribbons indicating 95 % confidence intervals (light-off control: grey; light-on control: black; light-off test: orange; light-on test: red). **F.** Time series of average locomotion speed (mm/s) for the same flies and conditions as in panel E. Plot design is otherwise identical, with means and 95 % confidence intervals shown for each group. **G.** Estimation plot of feed volume for fed male flies with Tdc2 activation. *Upper panel:* Observations of feed volume for individual flies of the indicated genotypes (driver control: black; responder control: grey; *Tdc2>Chr*: red). Grey bars and gapped lines indicate group means ± standard deviations. *Lower panel:* Effect size analysis using estimation statistics. *Lower left side:* Mean differences (Δ) between control and test flies within each light condition (light-off: control vs test; light-on: control vs test), shown as black dots and grey bars with 95 % confidence intervals (black lines) and bootstrap distribution (grey curves). *Lower right side:* Difference of mean differences (ΔΔ), shown as black dots and grey bars with 95 % confidence intervals (black lines) and bootstrap distribution (grey curves). When ΔΔ confidence intervals exclude zero, this indicates evidence for a light-dependent effect beyond baseline differences. *Key:* Genotype groups (Control, Test), light conditions, sample sizes (Ns), and effect sizes (Δ or ΔΔ) are provided in the accompanying table. **H.** Estimation plot of walking speed for fed male flies with Tdc2 activation. Same plot design as panel G, but for average locomotion speed. Full genotypes, light conditions, sample sizes (Ns), and statistics are provided in Supplementary Table 1.

**Figure S1.**
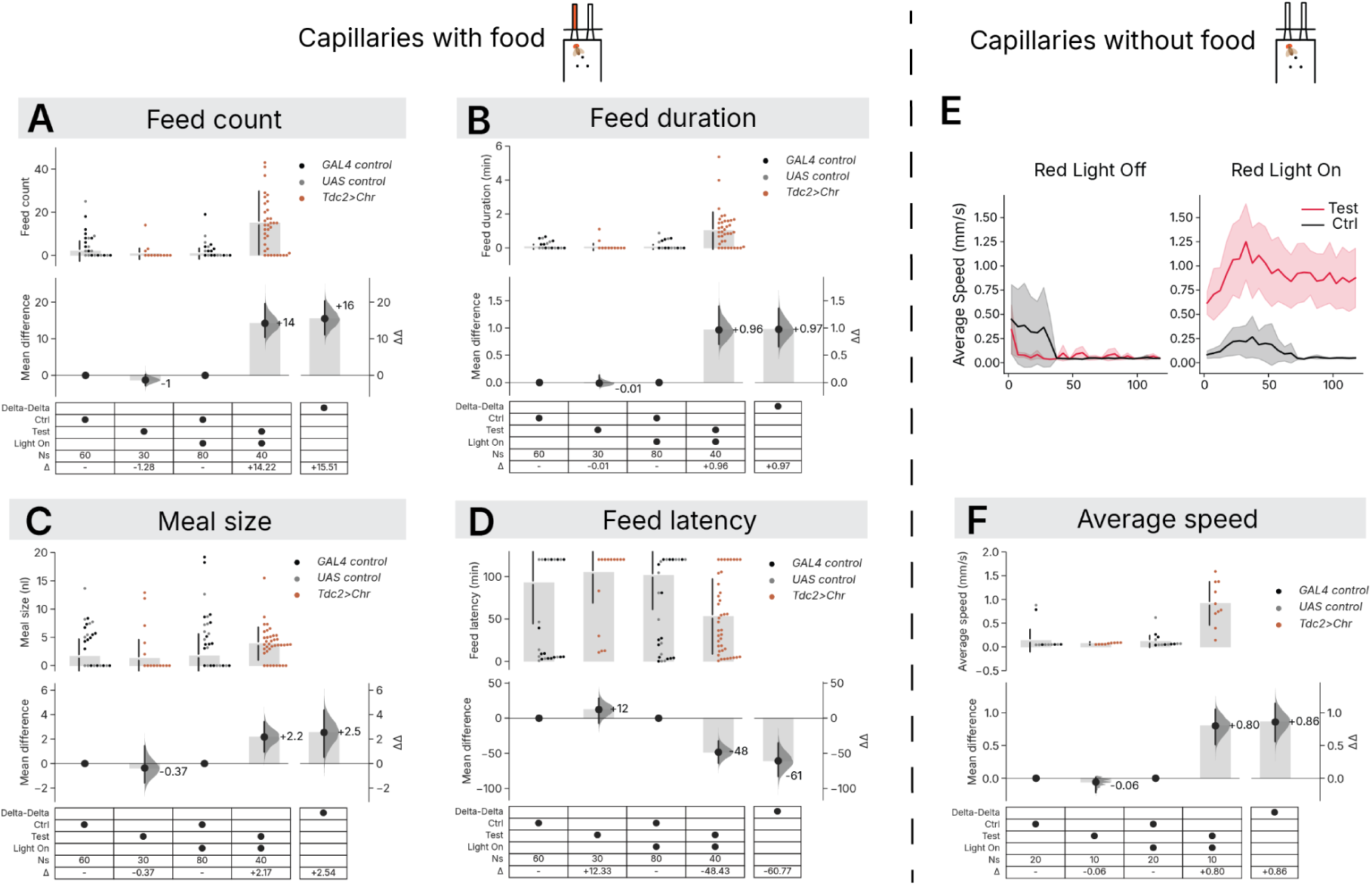
Effect of Tdc2-neuron activation on several metrics related to feeding. **A.** Estimation plot of feed count for fed male flies with Tdc2-neuron activation. **B.** Estimation plot of feed duration for fed male flies with Tdc2-neuron activation. **C.** Estimation plot of meal size for fed male flies with Tdc2-neuron activation. **D.** Estimation plot of feed latency for fed male flies with Tdc2-neuron activation. *Upper panel:* Observations of the respective metrics for individual flies of the indicated genotypes (driver control: black; responder control: grey; *Tdc2>Chr*: red). Grey bars and gapped lines indicate group means ± standard deviations. *Lower panel:* Effect size analysis using estimation statistics. *Lower left side:* Mean differences (Δ) between control and test flies within each light condition (light-off: control vs test; light-on: control vs test), shown as black dots and grey bars with 95 % confidence intervals (black lines) and bootstrap distribution (grey curves). *Lower right side:* Difference of mean differences (ΔΔ), shown as black dots and grey bars with 95 % confidence intervals (black lines) and bootstrap distribution (grey curves). When ΔΔ confidence intervals exclude zero, this indicates evidence for a light-dependent effect beyond baseline differences. *Key:* Genotype groups (Control, Test), light conditions, sample sizes (Ns), and effect sizes (Δ or ΔΔ) are provided in the accompanying table. **E.** Time series of average locomotion speed of fed male *Tdc2>Chr* flies and their respective controls during a 2-h assay using capillaries without food in red light-on and light-off conditions, with means and 95 % confidence intervals shown for each group. **F.** Estimation plot of average locomotion speed for the same experiment in panel E.

### Activation of two classes of specific octopaminergic neurons promote feeding

We hypothesized that the feeding phenotypes observed with broad Tdc2 neuron activation were mainly elicited by activity in specific OA-neuron classes. To test this, we examined two plausible candidates: the VPM4 neurons, previously implicated in feeding-associated behaviors, and VPM3 neurons, a neighboring MB-projecting neuron subset (labeled by split Gal4 line *XY101*, see Materials and Methods) (**Figure 2A-B**). Espresso analysis showed that both VPM3 and VPM4 independently promoted feeding in fed flies (**Figure 2C-D**). In addition, in 24-h starved flies, VPM4 activation further increased feed volume, suggesting these neurons either encode a signal that exceeds natural 24-h hunger or provide a hunger-independent feeding drive (**Figure S2B**). VPM3 activation effects varied across metabolic states. In fed males, VPM3 activation increased feed volume by ΔΔ = + 102 nl [95CI 63, 140] (**Figure 2C**), while in 24h-starved males, it increased feed volume by ΔΔ = + 89 nl [95CI -10, 184] (**Figure S2A**);, and the locomotion enhancement was absent in starved flies (**Figure S2C**). VPM4 activation effects also varied with metabolic state. Notably, the meal size effect showed opposite directions: in fed males Δg = 0.78 [95CI 0.18, 1.56] (ΔΔ = +8.23 nl [95CI 1.93, 16.45]) versus in starved males Δg = –0.68 [95CI –1.41, –0.02] (ΔΔ = –25.49 nl [95CI –53.11, -0.78]) (**Figure S2D**). These differential responses suggest that VPM3 and VPM4 interact with metabolic state in distinct ways.

**Figure 2.**
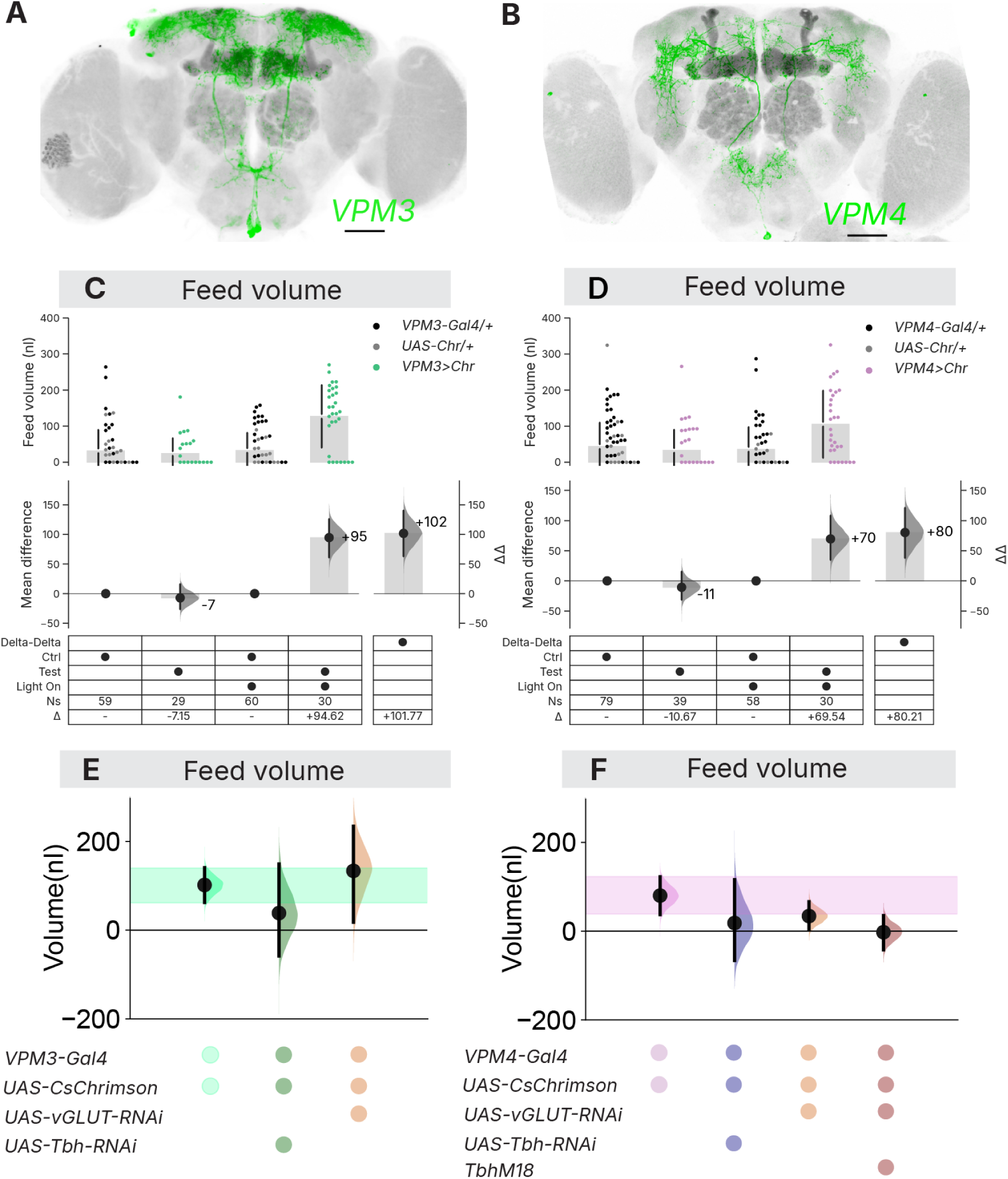
Two octopaminergic cell types, VPM3 and VPM4, promote food consumption. **A.** Expression of Venus, visualized by anti-GFP immunostaining, in VPM3 neurons using *XY101-Gal4*. Neuropils and the somata of Gal4-expressing cells, as revealed by anti-DLG (grey) and CsChrimson.mVenus (green), are shown, respectively. Scale bar = 50 µm. **B.** Expression of Venus in VPM4 neurons using *MB113-Gal4*. Plot elements as in panel A. **C.** Estimation plot of feed volume for fed male flies with VPM3 activation. *Upper panel:* Observations of feed volume for individual flies of the indicated genotypes (color coding: driver control: black, responder control: grey, VPM3 neuron activation: green). Grey bars and gapped lines indicate group means ± standard deviations. *Lower panel:* Effect size analysis using estimation statistics. *Lower left side:* Mean differences (Δ) between control and test flies within each light condition (light-off: control vs test; light-on: control vs test), shown as black dots and grey bars with 95 % confidence intervals (black lines) and bootstrap distribution (grey curves). *Lower right side:* Difference of mean differences (ΔΔ), shown as black dots and grey bars with 95 % confidence intervals (black lines) and bootstrap distribution (grey curves). When ΔΔ confidence intervals exclude zero, this indicates evidence for a light-dependent effect beyond baseline differences. **D.** Estimation plot of feed volume for fed male flies with VPM4 activation. Plot design is identical to panel C, except that the observed values of individual feed volumes are shown with purple dots. **E.** The VPM3-activation consumption increase requires octopamine. Forest plot showing differences of mean differences of feed volume (ΔΔ nl) in fed male flies under the following conditions: VPM3 activation alone; VPM3 activation with Tbh RNAi knockdown; or VPM3 activation with vGlut RNAi knockdown. Black dots represent ΔΔ effect sizes, with vertical black lines indicating their 95 % confidence intervals. Shaded distribution curves show the ΔΔ error. The green ribbon highlights the 95 % confidence interval for the VPM3 activation effect. The gridkey below the plot indicates experimental groups (VPM3>Chr, springgreen; VPM3>Chr, Tbh-RNAi, darkgreen; VPM3>Chr, vGlut-RNAi, chocolate). The estimation plots for each ΔΔ are displayed in Figure S4. **F.** The VPM4-activation consumption increase requires both octopamine and *VGlut1*. Forest plot showing differences of mean differences of feed volume (ΔΔ, nl) in fed male flies under the following conditions: VPM4 activation alone; VPM4 activation with *Tbh* RNAi knockdown; VPM4 activation with *VGlut1* RNAi knockdown; and VPM4 activation with both *Tbh^nM18^* background and *VGlut1* RNAi knockdown. Most elements are the same as in panel E. The purple ribbon highlights the 95 % confidence interval for the VPM4 activation effect. The gridkey indicates the experimental groups (VPM4>Chr, orchid; VPM4>Chr, Tbh-RNAi, navy; VPM4>Chr, vGlut-RNAi, chocolate; *Tbh^nM18^*, VPM4>Chr, vGlut-RNAi, maroon). The estimation plots for each ΔΔ are displayed in Figure S4. Genotype comparisons, light conditions, sample sizes (Ns), and effect sizes are provided in Supplementary Table 1.

Intriguingly, when we inhibited a broad range of OA neurons or specific OA populations using *Guillardia theta* Anion Channelrhodopsin 1, GtACR1 (Mohammad et al. 2017) or tetanus toxin, TeNT (Sweeney et al. 1995) in flies starved for 24 h, we did not see any consumption effect (**Figure S3A-D**). These results suggest that these OA neurons are not required for hunger-motivated feeding.

**Figure S2.**
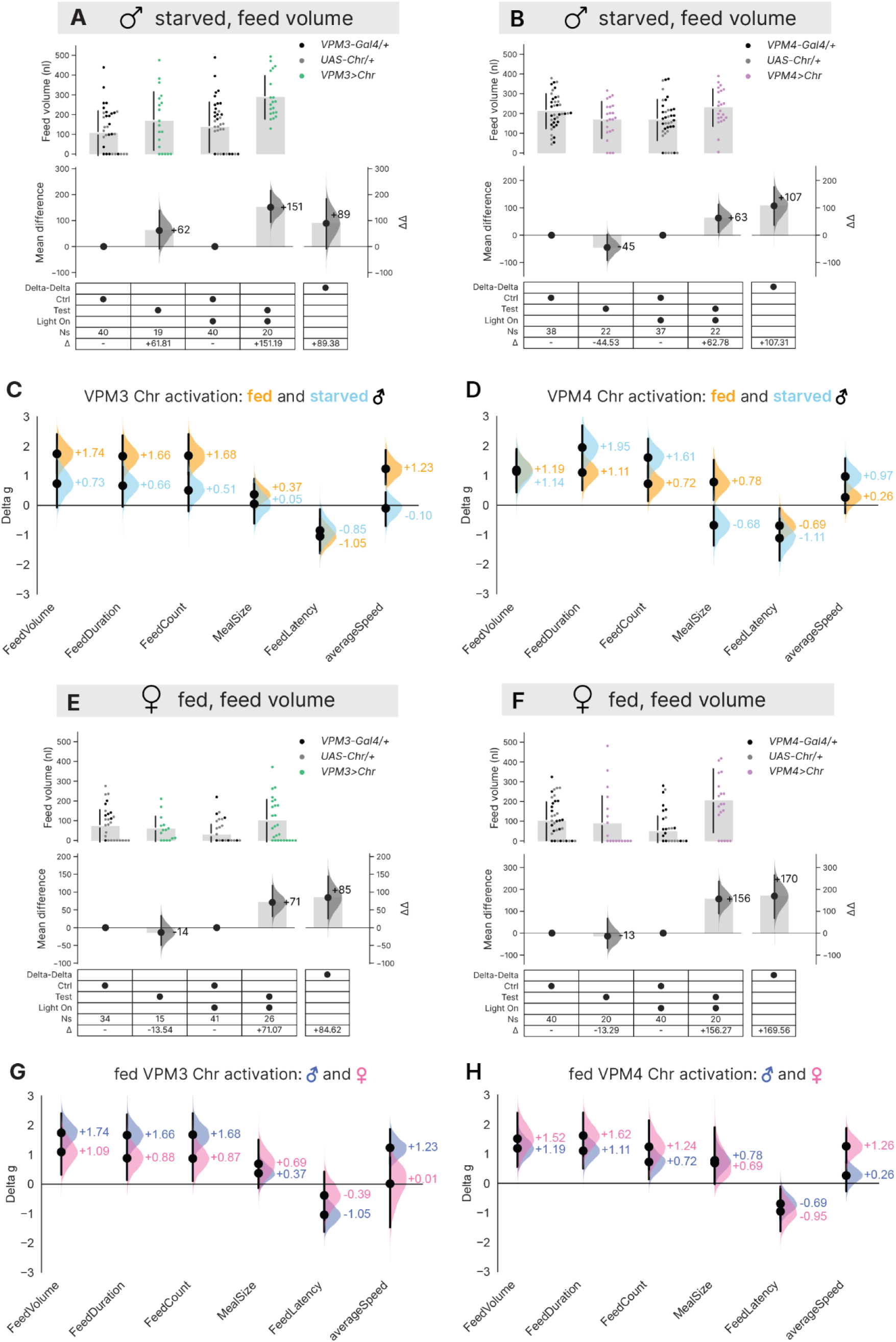
Activation of VPM3 or VPM4 increases feeding. **A.** Estimation plot of feed volume for 24-h starved (male) flies with VPM3 activation. **B.** Estimation plot of feed volume for 24-h starved (male) flies with VPM4 activation. **C.** Forest plot showing standardized effect sizes (Δg) for 6 feeding and locomotor metrics in Espresso of fed male (orange) and starved male (skyblue) flies with VPM3 activation. Black dots represent Δg, with vertical black lines indicating their 95 % confidence intervals. Shaded distribution curves show the Δg error. See Methods for the calculation of standardized effect sizes (Δg) from delta-delta (ΔΔ) values. **D.** Forest plot of standardized effect sizes (Δg) for 6 feeding and locomotor metrics in Espresso of fed male (orange) and starved male (skyblue) flies with VPM4 activation. Black dots represent Δg, with vertical black lines indicating their 95 % confidence intervals. Shaded distribution curves show the Δg error. **E.** Estimation plot of feed volume for female fed flies with VPM3 activation. **F.** Estimation plot of feed volume for female fed flies with VPM4 activation. *Upper panel:* Observations of feed volume for individual flies of the indicated genotypes. Grey bars and gapped lines indicate group means ± standard deviations. *Lower panel:* Effect size analysis using estimation statistics. *Lower left side:* Mean differences (Δ) between control and test flies within each light condition (light-off: control vs test; light-on: control vs test), shown as black dots and grey bars with 95 % confidence intervals (black lines) and bootstrap distribution (grey curves). *Lower right side:* Difference of mean differences (ΔΔ), shown as black dots and grey bars with 95 % confidence intervals (black lines) and bootstrap distribution (grey curves). When ΔΔ confidence intervals exclude zero, this indicates evidence for a light-dependent effect beyond baseline differences. **G.** Forest plot of standardized effect sizes (Δ*g* for 6 feeding and locomotor metrics in Espresso of fed male (royal blue) and fed female (pink) flies with VPM3 activation. Black dots represent Δg, with vertical black lines indicating their 95 % confidence intervals. Shaded distribution curves show the Δg error. **H.** Forest plot of standardized effect sizes (Δ*g*) for 6 feeding and locomotor metrics in Espresso of fed male (royal blue) and fed female (pink) flies with VPM4 activation. Black dots represent Δg, with vertical black lines indicating their 95 % confidence intervals. Shaded distribution curves show the Δg error. Genotype comparisons, light conditions, sample sizes (Ns), and effect sizes are provided in Supplementary Table 1.

### VPM-neuron activation promotes feeding behaviors via octopamine release

VPM4 has been previously shown to co-transmit OA and glutamate (Sherer et al. 2020). In this study, RNA-interference (RNAi) knockdown of the OA-synthase *Tbh* in flies expressing Chr in VPM3 or VPM4 neurons abolished all feeding phenotypes associated with neuronal activation, indicating that the VPM-activation effects were strongly OA-dependent (**Figure 2E-F, Figure S4, Figure S5**). Additionally, knockdown of *vesicular glutamate transporter 1* (*VGlut1*) in VPM4 neurons attenuated the effect of activation on feed volume from +80 nl to +34 nl and abolished the effects on meal size and latency, while feed count and duration effects remained unchanged (**Figure 2F, Figure S5G-L**). In flies carrying both *VGlut1* RNAi and a *Tbh^nM18^* mutation, VPM4 activation exhibited no feeding phenotypes, consistent with the effect observed in Tbh knockdown alone. (**Figure 2F, Figure S5G-L**), supporting the idea that glutamate signaling plays a minor role in mediating VPM4 feeding behaviors. These data show that the feeding effects of VPM3 and VPM4 neurons are primarily mediated via OA transmission.

### VPM-activity effects have limited sexual dimorphism

Octopaminergic neurons can have sexually dimorphic behavioral effects, including courtship, aggression, and post-mating responses (Certel et al. 2007; Machado et al. 2017; Rezával et al. 2014). VPM3 neurons show sex-dimorphic effects on sleep regulation and male courtship (Reyes et al. 2025). To investigate potential sex differences in feeding, we also examined VPM3 and VPM4 activation in previously fed female flies (**Figure S2E-H**). Both neuron types promoted feeding in sated females, though VPM3 activation reduced feed latency in fed males with Δg = –1.05 [95CI –1.61, –0.44] (ΔΔ = –47.26 min [95CI –72.56, –19.77]) but had no latency effect in females (**Figure S2G**). The locomotor effects showed more sexual dimorphism: VPM3 activation increased average walking speed in fed males but not in females, whereas VPM4 activation increased speed in fed females but not in males (**Figure S2G-H**). ACR1-mediated inhibition of VPM3 and VPM4 had mostly negligible effects in both sexes, and no effect on feed volume (**Figure S3E-F**). There were a few exceptions: VPM4 inhibition elicited a reduction in feed frequency in starved females and, in both sexes, increased meal size and decreased walking speed (**Figure S3H**). These findings indicate that VPM-neuron activity promotes feeding in both sexes and the optogenetic effects have relatively modest sexual dimorphism—mostly in locomotion—while the instructive feeding drive is similar in both sexes.

**Figure S3.**
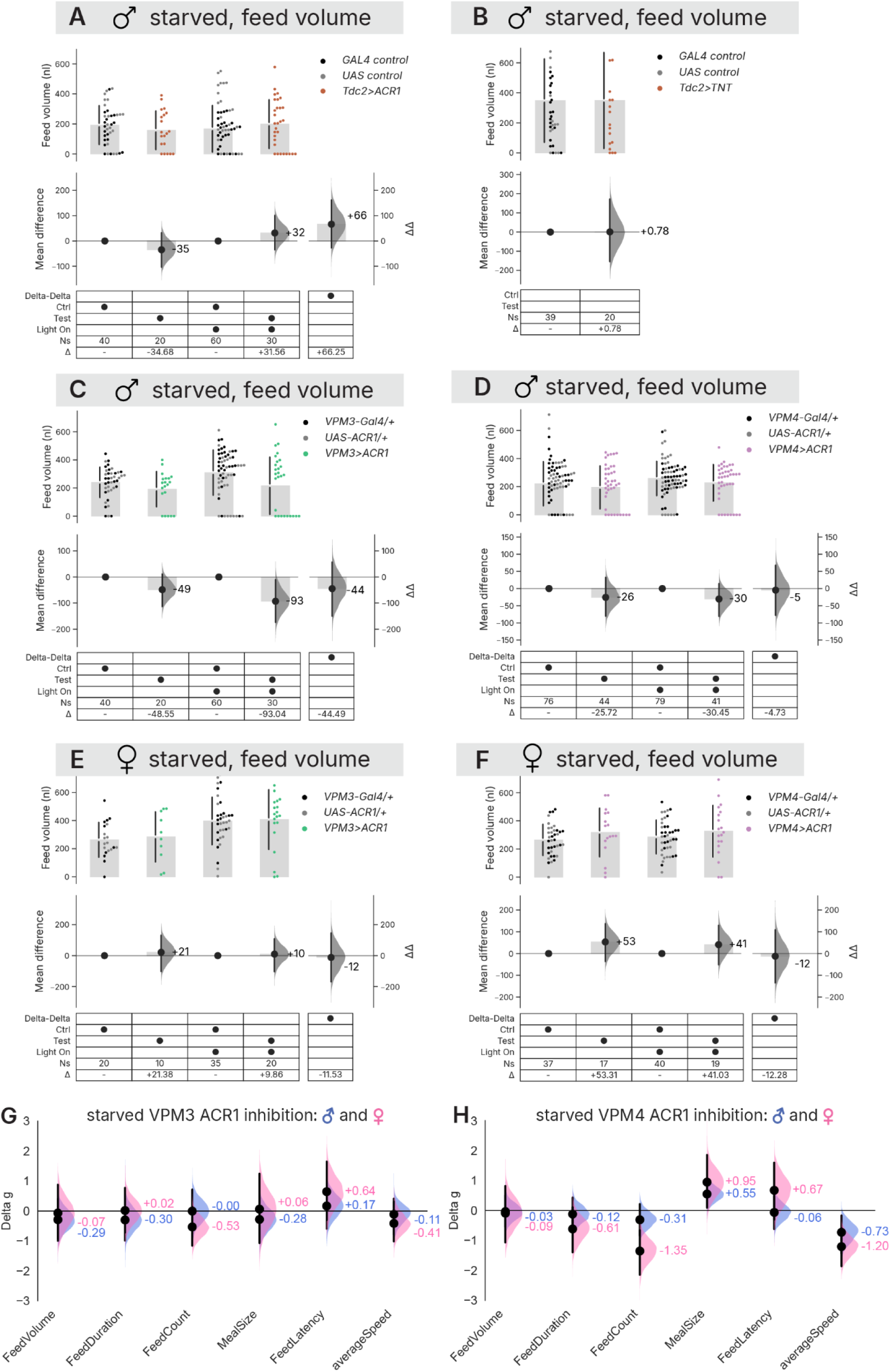
Inhibiting OA neurons in hungry flies has little effect on feed volume. **A.** Estimation plot of feed volume for 24-h starved male flies with Tdc2-cell inhibition. **B.** Estimation plot of feed volume of 24-h starved male flies expressing tetanus toxin (TeNT) in *Tdc2-Gal4* cells. *Upper panel:* Observations of feed volume for individual flies of the indicated genotypes (color coding: driver control: black, responder control: grey, Tdc2 neuron synaptic output block: orangered). Grey bars and gapped lines indicate group means ± standard deviations. *Lower panel:* mean differences (Δ) between controls and test (*Tdc2>TeNT*) flies, shown as black dots and grey bars with 95 % confidence intervals (black lines) and bootstrap distribution (grey curves). **C.** Estimation plot of feed volume for 24-h starved male flies with VPM3 inhibition. **D.** Volume for 24-h starved male flies with VPM4 inhibition. **E.** Volume for 24-h starved female flies with VPM3 inhibition. **F.** Volume for 24-h starved female flies with VPM4 inhibition. *Upper panel:* Observations of feed volume for individual flies of the indicated genotypes. Grey bars and gapped lines indicate group means ± standard deviations. *Lower panel:* Effect size analysis using estimation statistics. *Lower left side:* Mean differences (Δ) between control and test flies within each light condition (light-off: control vs test; light-on: control vs test), shown as black dots and grey bars with 95 % confidence intervals (black lines) and bootstrap distribution (grey curves). *Lower right side:* Difference of mean differences (ΔΔ), shown as black dots and grey bars with 95 % confidence intervals (black lines) and bootstrap distribution (grey curves). When ΔΔ confidence intervals exclude zero, this indicates evidence for a light-dependent effect beyond baseline differences. **G.** Forest plot of standardized effect sizes (Δg) for 6 feeding and locomotor metrics in Espresso of 24-h starved male (royal blue) and 24-h starved female (pink) flies with VPM3 inhibition. Black dots represent Δg, with vertical black lines indicating their 95 % confidence intervals. Shaded distribution curves show the Δg error. **H.** Forest plot of standardized effect sizes (Δg) for 6 feeding and locomotor metrics in Espresso of 24-h starved male (royal blue) and 24-h starved female (pink) flies with VPM4 inhibition. Black dots represent Δg, with vertical black lines indicating their 95 % confidence intervals. Shaded distribution curves show the Δg error. Genotype comparisons, light conditions, sample sizes (Ns), and effect sizes are provided in Supplementary Table 1.

**Figure S4.**
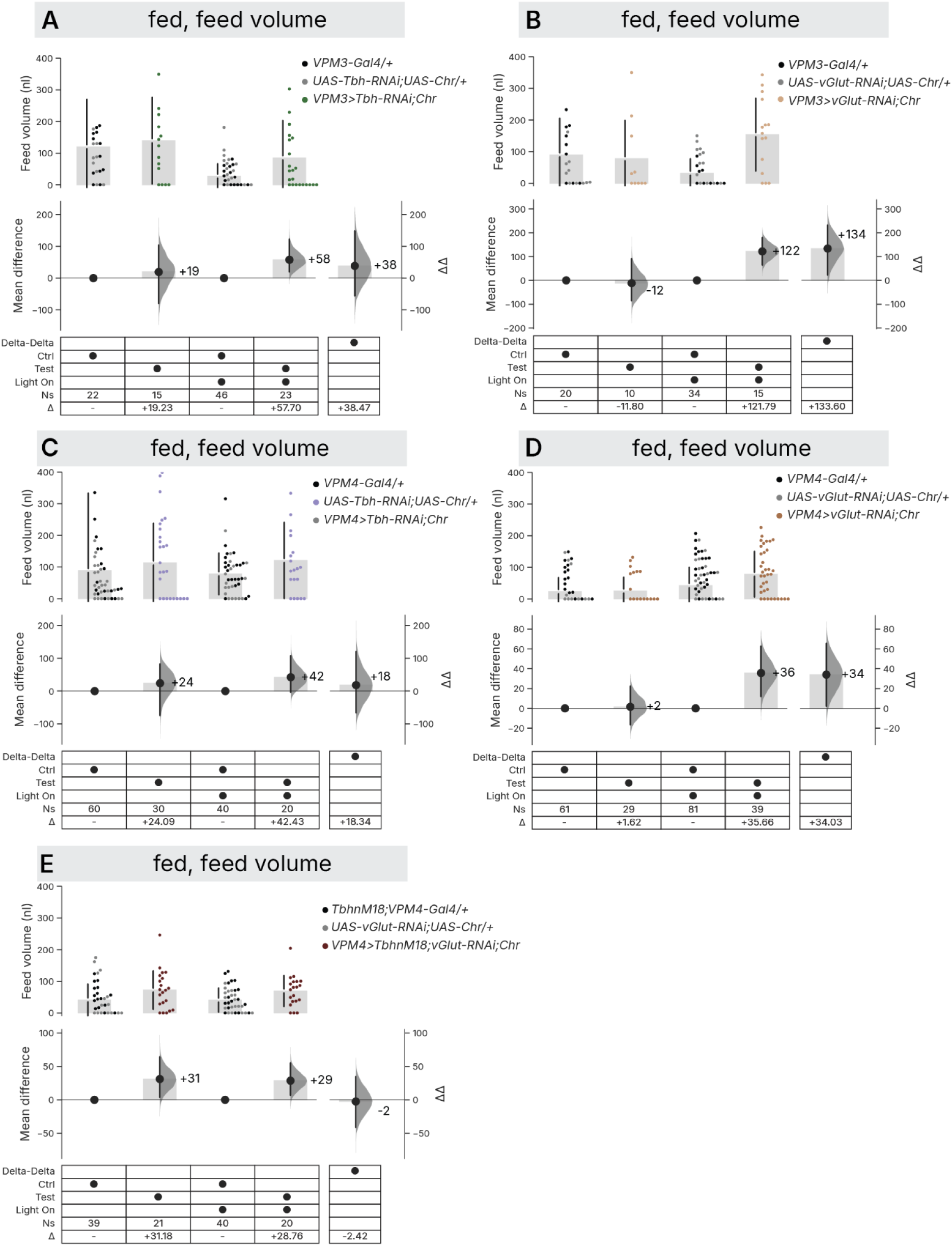
feed volume effects from VPM activation are mainly OA-dependent. **A.** Estimation plot of feed volume and effect sizes for VPM3 activation with *Tbh* RNAi knockdown in fed male flies. **B.** Volume effect with VPM3 activation with *VGlut1* RNAi knockdown fed male flies. **C.** Volume effect for VPM4 activation with *Tbh* RNAi knockdown in fed male flies. **D.** Volume effect for VPM4 activation with *VGlut1* RNAi knockdown in fed male flies. **E.** Volume effect for VPM4 activation with both *Tbh^nM18^* background and *VGlut1* RNAi knockdown in fed male flies. *Upper panel:* Observations of feed volume for individual flies of the indicated genotypes. Grey bars and gapped lines indicate group means ± standard deviations. *Lower panel:* Effect size analysis using estimation statistics. *Lower left side:* Mean differences (Δ) between control and test flies within each light condition (light-off: control vs test; light-on: control vs test), shown as black dots and grey bars with 95 % confidence intervals (black lines) and bootstrap distribution (grey curves). *Lower right side:* Difference of mean differences (ΔΔ), shown as black dots and grey bars with 95 % confidence intervals (black lines) and bootstrap distribution (grey curves). When ΔΔ confidence intervals exclude zero, this indicates evidence for a light-dependent effect beyond baseline differences. Genotype comparisons, light conditions, sample sizes (Ns), and effect sizes are provided in Supplementary Table 1.

**Figure S5.**
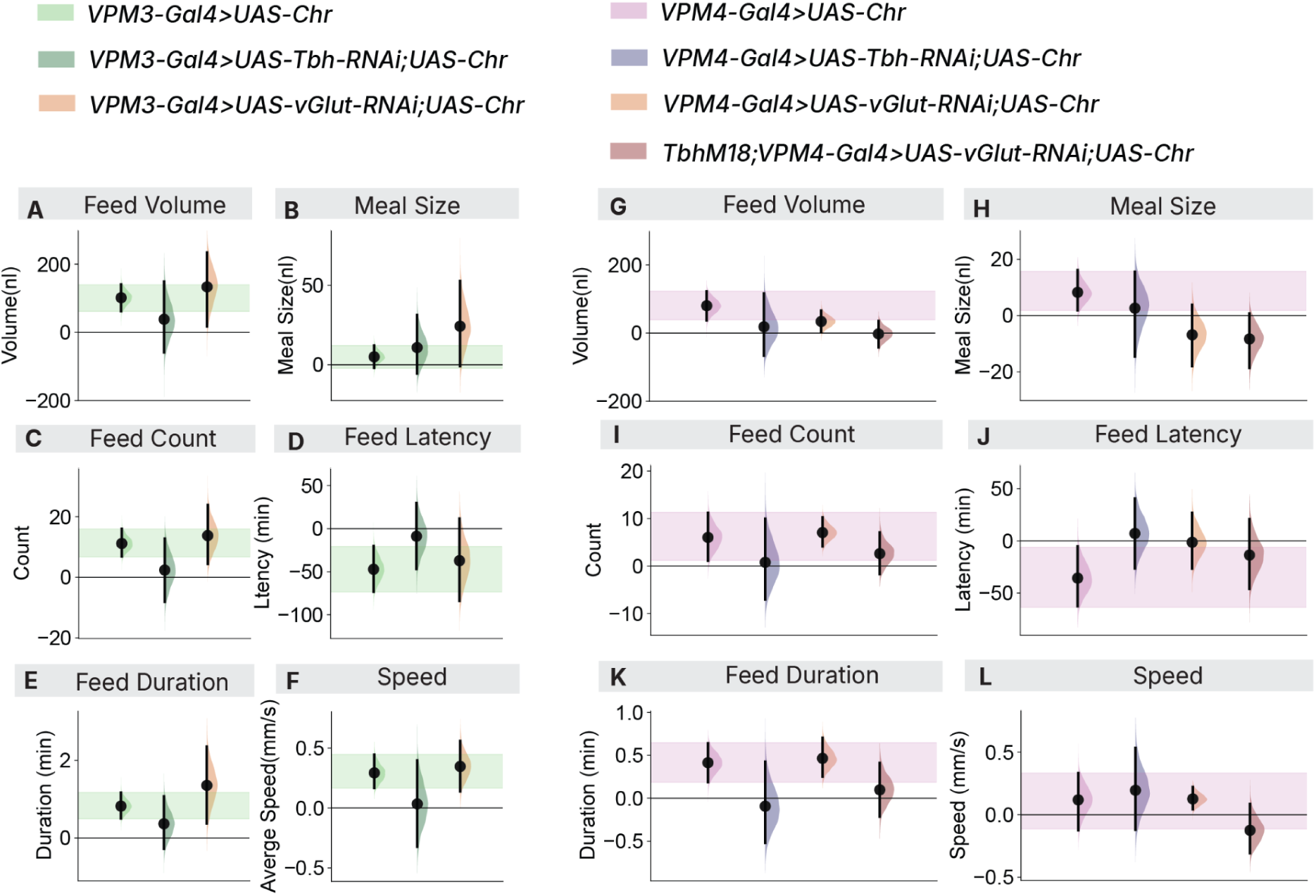
VPM effects on various feeding-related metrics are mainly OA-dependent. **A.** Forest plot showing differences of mean differences of feed volume (ΔΔ nl) in fed male flies under the following conditions: VPM3 activation alone; VPM3 activation with Tbh RNAi knockdown; or VPM3 activation with vGlut RNAi knockdown. Black dots represent ΔΔ effect sizes, with vertical black lines indicating their 95 % confidence intervals. Shaded distribution curves show the ΔΔ error. The green ribbon highlights the 95 % confidence interval for the VPM3 activation effect. The gridkey below the plot indicates experimental groups (VPM3>Chr, springgreen; VPM3>Chr, Tbh-RNAi, darkgreen; VPM3>Chr, vGlut-RNAi, chocolate). **B.** Differences of meal size (ΔΔ nl) in fed male flies under the following conditions: VPM3 activation alone; VPM3 activation with Tbh RNAi knockdown; or VPM3 activation with vGlut RNAi knockdown (for the same flies and conditions as in panel A). **C.** Differences of feed count (ΔΔ) in fed male flies under the following conditions: VPM3 activation alone; VPM3 activation with Tbh RNAi knockdown; or VPM3 activation with vGlut RNAi knockdown (for the same flies and conditions as in panel A). **D.** Differences of feed latency (ΔΔ min) in fed male flies under the following conditions: VPM3 activation alone; VPM3 activation with Tbh RNAi knockdown; or VPM3 activation with vGlut RNAi knockdown (for the same flies and conditions as in panel A). **E.** Differences of feed duration (ΔΔ min) in fed male flies under the following conditions: VPM3 activation alone; VPM3 activation with Tbh RNAi knockdown; or VPM3 activation with vGlut RNAi knockdown (for the same flies and conditions as in panel A). **F.** Differences of locomotion speed (ΔΔ mm/s) in fed male flies under the following conditions: VPM3 activation alone; VPM3 activation with Tbh RNAi knockdown; or VPM3 activation with vGlut RNAi knockdown (for the same flies and conditions as in panel A). **G.** Differences of feed volume (ΔΔ, nl) in fed male flies under the following conditions: VPM4 activation alone; VPM4 activation with *Tbh* RNAi knockdown; VPM4 activation with *VGlut1* RNAi knockdown; and VPM4 activation with both *Tbh^nM18^* background and *VGlut1* RNAi knockdown. Most elements are the same as in panel A. The purple ribbon highlights the 95 % confidence interval for the VPM4 activation effect. The gridkey indicates the experimental groups (VPM4>Chr, orchid; VPM4>Chr, Tbh-RNAi, navy; VPM4>Chr, vGlut-RNAi, chocolate; *Tbh^nM18^*, VPM4>Chr, vGlut-RNAi, maroon). **H.** Differences of meal size (ΔΔ nl) in fed male flies under the following conditions: VPM4 activation alone; VPM4 activation with *Tbh* RNAi knockdown; VPM4 activation with *VGlut1* RNAi knockdown; and VPM4 activation with both *Tbh^nM18^* background and *VGlut1* RNAi knockdown (for the same flies and conditions as in panel G). **I.** Differences of feed count (ΔΔ) in fed male flies under the following conditions: VPM4 activation alone; VPM4 activation with *Tbh* RNAi knockdown; VPM4 activation with *VGlut1* RNAi knockdown; and VPM4 activation with both *Tbh^nM18^* background and *VGlut1* RNAi knockdown (for the same flies and conditions as in panel G). **J.** Differences of feed latency (ΔΔ min) in fed male flies under the following conditions: VPM4 activation alone; VPM4 activation with *Tbh* RNAi knockdown; VPM4 activation with *VGlut1* RNAi knockdown; and VPM4 activation with both *Tbh^nM18^* background and *VGlut1* RNAi knockdown (for the same flies and conditions as in panel G). **K.** Differences of feed duration (ΔΔ min) in fed male flies under the following conditions: VPM4 activation alone; VPM4 activation with *Tbh* RNAi knockdown; VPM4 activation with *VGlut1* RNAi knockdown; and VPM4 activation with both *Tbh^nM18^* background and *VGlut1* RNAi knockdown (for the same flies and conditions as in panel G). **L.** Differences of locomotion speed (ΔΔ mm/s) in fed male flies under the following conditions: VPM4 activation alone; VPM4 activation with *Tbh* RNAi knockdown; VPM4 activation with *VGlut1* RNAi knockdown; and VPM4 activation with both *Tbh^nM18^* background and *VGlut1* RNAi knockdown (for the same flies and conditions as in panel G).

### A major postsynaptic target of VPM cells in the mushroom body is MBON11

We then sought to identify the downstream circuits through which VPM neurons influence different feeding phenotypes. Published adult fly brain connectome data show that both VPM3 and VPM4 have synaptic connectivity with MBON11 in the peduncles of the MBs (VPM3: 80 connections, VPM4: 83 connections) (**Figure 3A-B**). No other OA neurons were found to have direct connections with MBON11 (Sayin et al. 2019; Scheffer et al. 2020; Plaza et al. 2022). To corroborate this connectivity, we used GFP reconstitution across synaptic partners (GRASP) (Feinberg et al. 2008) to detect interactions between the Tdc2+ neurons (*Tdc2-LexA*) and MBON11 neurons (*MB112C-Gal4*); this experiment revealed anatomical contact between octopaminergic (likely VPM) neurons and MBON11 neurons in the MB lobes, supporting the presence of a synaptic connection (**Figure 3C**).

**Figure 3.**
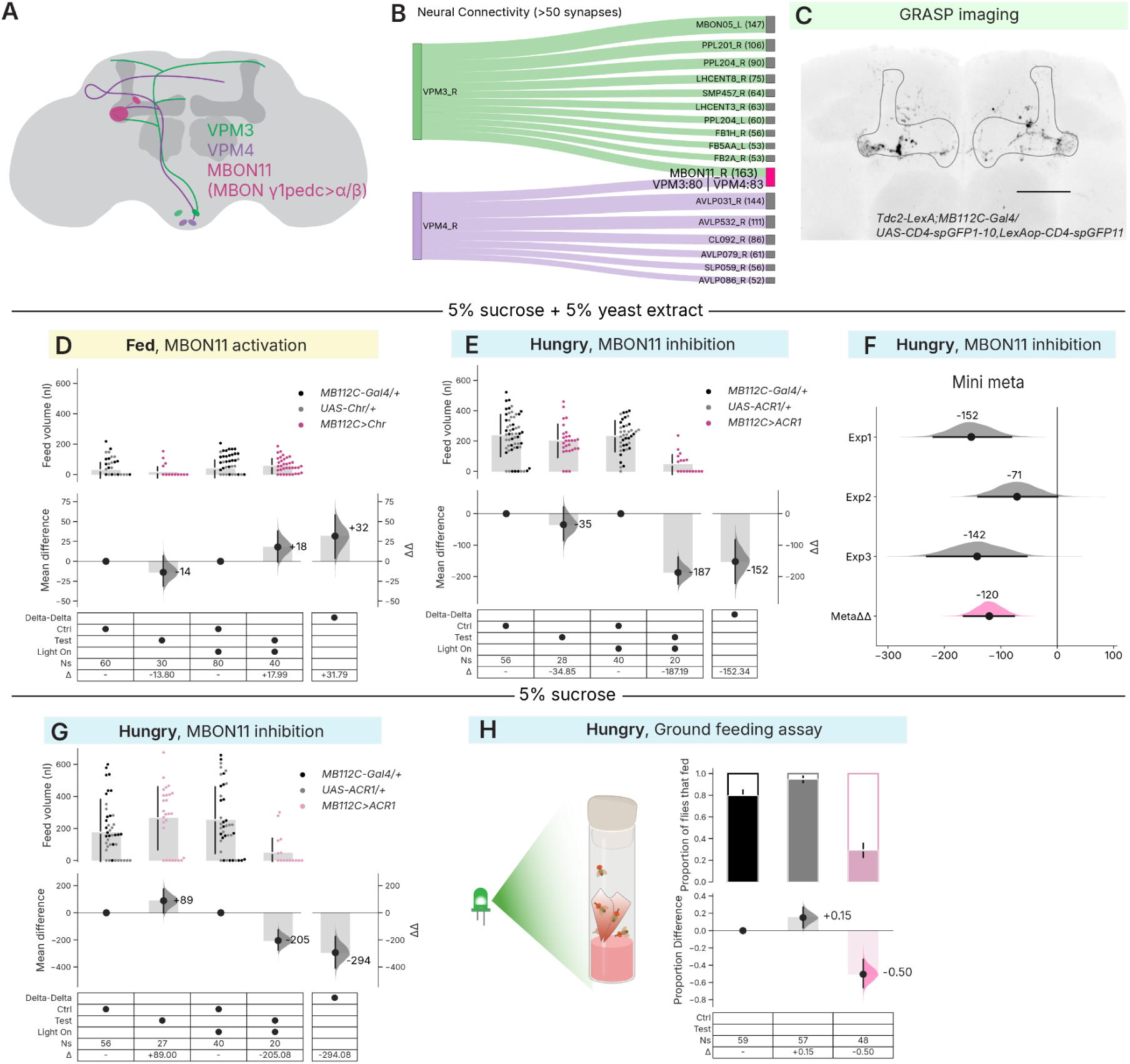
The VPM-postsynaptic partner MBON11 regulates feeding. **A.** An illustration of the projection patterns of VPM3, VPM4, and MBON11. **B.** Postsynaptic connections to VPM3 and VPM4 (Synapse>50). Sankey plot shows the cell type of the downstream individual neurons that have strong postsynaptic connections (Synapse>50) with VPM3 and VPM4, with the width of the link proportional to the count of the connection. The number of connections is indicated inside the brackets next to the cell types. Cell types from the left (L) or right (R) hemisphere were designated with an underscore prefix. **C.** Native GRASP signals between Tdc2 and MBON11 (MB112C) neurons. The outline of the mushroom-body lobes are traced in black. The scale bar is 50 µm. **D.** Estimation plot of feed volume (5 % sucrose +5 % yeast extract) for fed male flies with MBON11 activation. **E.** Estimation plot of feed volume (5 % sucrose +5 % yeast extract) for 24-h starved male flies with MBON11 inhibition. **F.** Difference of mean differences (ΔΔ) of feed volume from three independent experiments of MBON11 inhibition in 24-h starved male flies and their combined meta-analysis effect size. **G.** Estimation plot of feed volume (5 % sucrose) for 24-h starved male flies with MBON11 inhibition. *Upper panel:* Observations of feed volume for individual flies of the indicated genotypes (color coding: driver control: black, responder control: grey, MBON11 neuron manipulation: pink). Grey bars and gapped lines indicate group means ± standard deviations. *Lower panel:* Effect size analysis using estimation statistics. *Lower left side:* Mean differences (Δ) between control and test flies within each light condition (light-off: control vs test; light-on: control vs test), shown as black dots and grey bars with 95 % confidence intervals (black lines) and bootstrap distribution (grey curves). *Lower right side:* Difference of mean differences (ΔΔ), shown as black dots and grey bars with 95 % confidence intervals (black lines) and bootstrap distribution (grey curves). When ΔΔ confidence intervals exclude zero, this indicates evidence for a light-dependent effect beyond baseline differences. **H**. Proportion of 24-h starved MBON11>ACR1 male flies and their respective driver and responder controls that fed on 5 % sucrose (had red color in abdomen) during a 2-h ground feeding assay. *Upper panel:* bar plots colored portion represents the proportion of flies that fed on 5 % sucrose while the white portion represents the proportion of flies that did not feed on 5 % sucrose (of indicated genotypes); the gapped lines represent the means with standard deviation. *Lower panel:* mean differences (Δ) between responder control and driver control, or *MBON11>ACR1* test flies and driver control, with 95CI (black bar) and distribution (grey curve). Genotype comparisons, light conditions, sample sizes (Ns), and effect sizes are provided in Supplementary Table 1.

**Figure S6.**
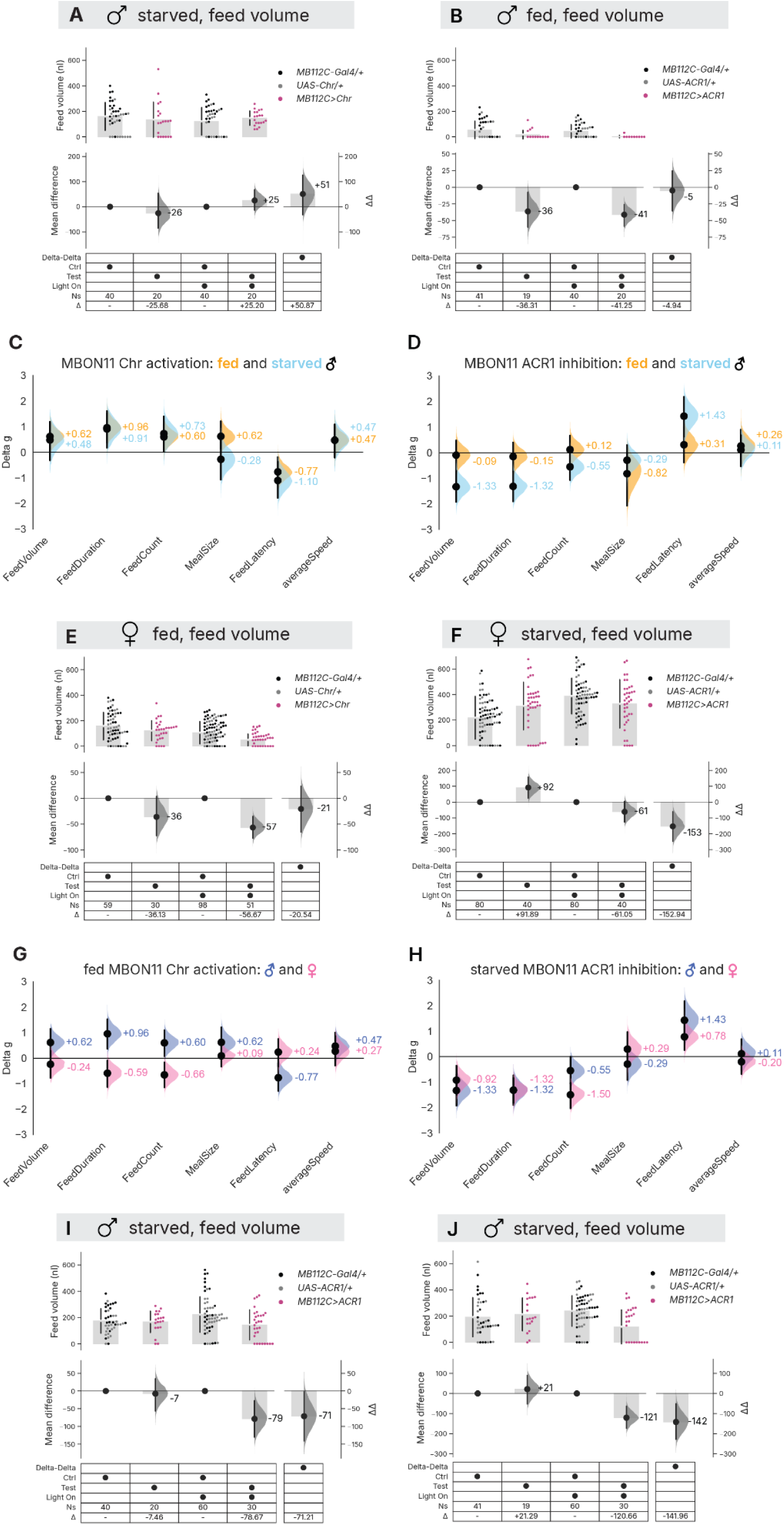
In fed and starved flies, MBON11 activation and inhibition have bidirectional effects on several feeding features and show sexual dimorphism. **A.** Estimation plot of feed volume for 24-h starved male flies with MBON11 activation. **B.** Estimation plot of feed volume for fed male flies with MBON11 inhibition. **C.** Forest plot showing standardized effect sizes (Δg) for 6 feeding and locomotor metrics in Espresso of fed male (orange) and starved male (skyblue) flies with MBON11 activation. Black dots represent Δg, with vertical black lines indicating their 95 % confidence intervals. Shaded distribution curves show the Δg error. **D.** Forest plot of standardized effect sizes (Δg) for 6 feeding and locomotor metrics in Espresso of fed male (orange) and starved male (skyblue) flies with MBON11 activation. Black dots represent Δg, with vertical black lines indicating their 95 % confidence intervals. Shaded distribution curves show the Δg error. **E.** Estimation plot of feed volume for fed female flies with MBON11 activation. **F.** Estimation plot of feed volume for 24-h starved female flies with MBON11 inhibition. **G.** Forest plot of standardized effect sizes (Δg) for 6 feeding and locomotor metrics in Espresso of fed male (royal blue) and fed female (pink) flies with MBON11 activation. Black dots represent Δg, with vertical black lines indicating their 95 % confidence intervals. Shaded distribution curves show the Δg error. **H.** Forest plot of standardized effect sizes (Δg) for 6 feeding and locomotor metrics in Espresso of 24-h starved male (royal blue) and 24-h starved female (pink) flies with MBON11 inhibition. Black dots represent Δg, with vertical black lines indicating their 95 % confidence intervals. Shaded distribution curves show the Δg error. **I.** Estimation plots of feed volume for 24-h starved male flies with MBON11 inhibition in an independent experiment 2, shown in mini meta plot in Figure 3F. **J.** Estimation plots of feed volume for 24-h starved male flies with MBON11 inhibition in an independent experiment 3, shown in mini meta plot in Figure 3F. *Upper panel:* Observations of feed volume for individual flies of the indicated genotypes. Grey bars and gapped lines indicate group means ± standard deviations. *Lower panel:* Effect size analysis using estimation statistics. *Lower left side:* Mean differences (Δ) between control and test flies within each light condition (light-off: control vs test; light-on: control vs test), shown as black dots and grey bars with 95 % confidence intervals (black lines) and bootstrap distribution (grey curves). *Lower right side:* Difference of mean differences (ΔΔ), shown as black dots and grey bars with 95 % confidence intervals (black lines) and bootstrap distribution (grey curves). When ΔΔ confidence intervals exclude zero, this indicates evidence for a light-dependent effect beyond baseline differences. Genotype comparisons, light conditions, sample sizes (Ns), and effect sizes are provided in Supplementary Table 1.

### Instructive and required: MBON11 activity has two-way control over feeding

Previous studies have suggested a role for MBON11 in food seeking, but we hypothesized this could be secondary to a more fundamental function in hunger-state control. We therefore directly tested whether MBON11 activity influences actual food consumption and feeding behavior. In previously *ab libitum* fed flies, MBON11 activation drove hunger-like behavior: it reduced feed latency and increased feed volume, feed count, feed duration, and meal size (**Figure 3D, Figure S6C**). Conversely, in 24-h starved flies, inhibition of MBON11 activity using GtACR1 actuation resulted in satiety-like behavior: delayed feed latency and drastically reduced food intake, feed count, and feeding duration (**Figure 3E-F, Figure S6D**). Although the flies remained lower in the chamber, MBON11 inhibition caused no other measurable deficits in locomotion or climbing ability (**Figure S6D, Figure S7C, F**). The feeding reduction effect of GtACR1 is reproducible: the finding was replicated across two additional iterations, and a meta-analysis of all three experiments yielded a combined effect size (meta ΔΔ) of –120 nl (**Figure 3F**). Inhibition with GtACR1 reduced food consumption by ∼70 % (**Figure 3E**), supporting the idea that MBON11 activity is required for a large fraction of hunger-motivated feeding.

We also tested optogenetic interventions under opposite nutritional conditions. In starved flies, MBON11 activation increased feed duration and reduced feed latency (**Figure S6A, C**). In fed flies, however, MBON11 inhibition had no detectable effect to further increase satiety (**Figure S6B, F**). Together, these findings highlight that MBON11 activity has a bidirectional influence: high activity drives increased feeding, while low activity elicits reduced feeding, specifically hunger-motivated feeding. This differentiates its function from the VPMs, which show no inhibition effect, so are not required for natural hunger-driven feeding, despite their ability to promote feeding when activated. Together, these results support that MBON11 activity in hungry flies has a permissive role in hunger-motivated feeding, and MBON11 activity levels are both required for and instructive to feeding.

### MBON11 feeding effects are not dependent on olfaction or climbing ability

Previous studies have implicated MBON11 neurons in the modulation of olfactory behaviors (Perisse et al. 2016; Tsao et al. 2018a; Matheson et al. 2022). We thus hypothesized that the feeding effects due to MBON11 activation or inhibition could arise from a strong dependency upon changes in odor attraction. To test this hypothesis, we presented flies with capillaries with sucrose without yeast extract (**Figure 3G**). In hungry flies presented with this minimally odorous sugar-only food, MBON11 inhibition still caused a robust suppression of food consumption (ΔΔ = −294 nl [95CI −408, −174]). As sugar water emits little odor, this result is consistent with the idea that MBON11 feeding effects are not strongly dependent on olfactory function.

In Espresso, MBON11-inhibited flies tended to remain at the bottom of the chamber (**Figure S7C, F**). To examine the potential confound of MBON11 effects on climbing, we implemented an assay with accessible food. Both *MB112C>ACR1* and control flies were placed in vials containing red dye- and sucrose-soaked tissues embedded in an agarose floor; flies with red liquid in their abdomen were counted after 2 h (**Figure 3H**). Consistent with the Espresso findings, approximately 60 % fewer flies fed when their MBON11 neurons were inhibited, confirming that reduced feeding is not a consequence of their inability to climb to the ceiling food source (**Figure 3H**). While MBON11 neuronal interventions clearly have pleiotropic effects on feeding and locomotion patterns, these experiments demonstrate that, rather than merely reflecting an epiphenomenal effect on climbing ability, MBON11 activity has authentic control over feeding behavior per se, including eating.

**Figure S7.**
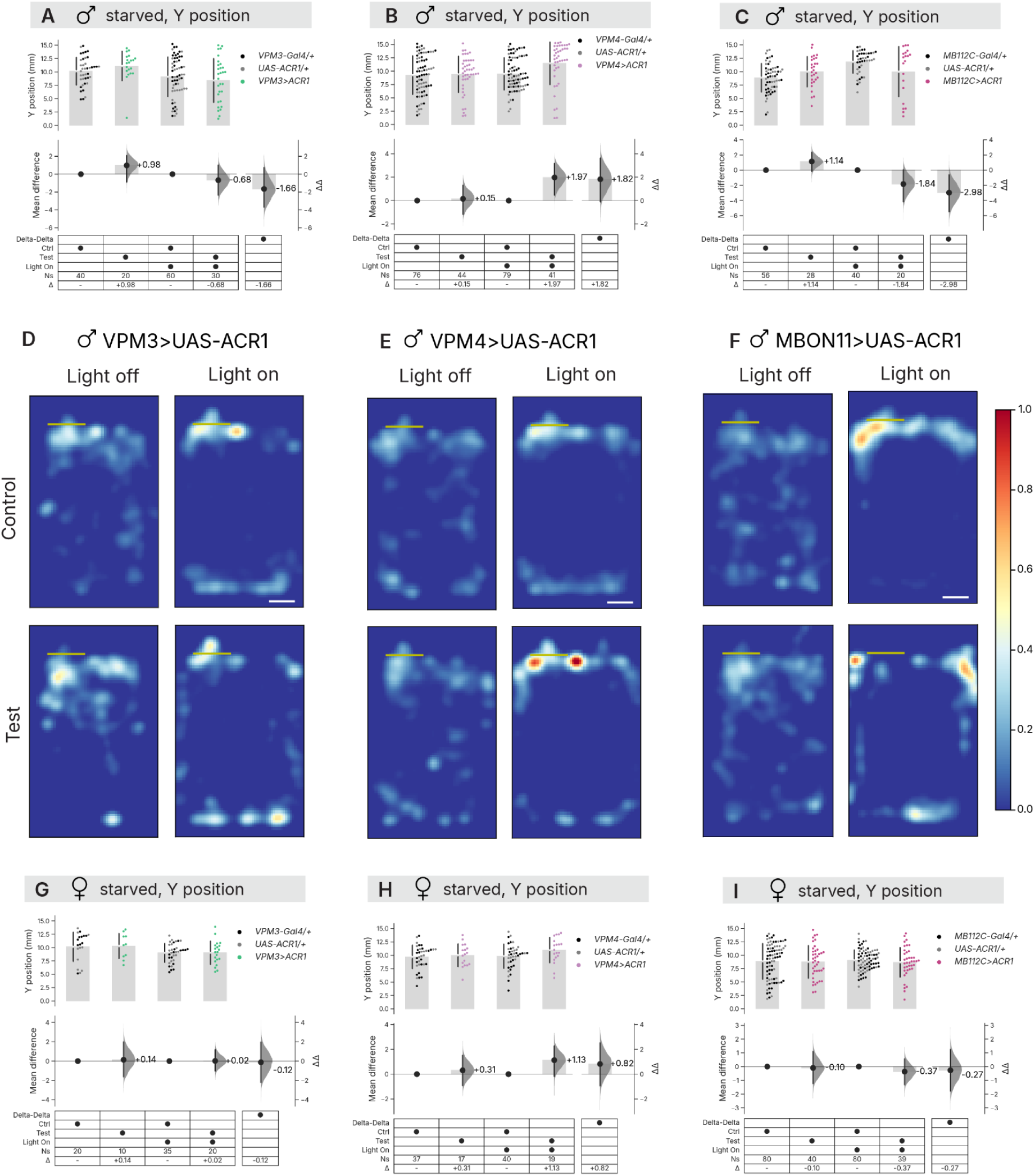
MBON11 inhibition reduces average elevation in hungry male flies. **A.** Estimation plots of elevation (Y position) for 24-h starved male flies with VPM3 opto-inhibition. **B.** Estimation plots of elevation (Y position) for 24-h starved male flies with VPM4 opto-inhibition. **C.** Estimation plots of elevation (Y position) for 24-h starved male flies with MBON11 opto-inhibition. **D.** Heatmaps showing the z score of time that 24-h starved male flies (of indicated groups and conditions) spent at different locations in the Espresso fly chambers with VPM3 opto-inhibition. **E.** Heatmaps showing the z score of time that 24-h starved male flies (of indicated groups and conditions) spent at different locations in the Espresso fly chambers with VPM4 opto-inhibition. **F.** Heatmaps showing the z score of time that 24-h starved male flies (of indicated groups and conditions) spent at different locations in the Espresso fly chambers with MBON11 opto-inhibition. **G.** Estimation plot of elevation (Y position) for 24-h starved female flies with VPM3 opto-inhibition. **H.** Estimation plot of elevation (Y position) for 24-h starved female flies with VPM4 opto-inhibition. **I.** Estimation plot of elevation (Y position) for 24-h starved female flies with MBON11 opto-inhibition.*Upper panel:* Observations of elevation (Y position) for individual flies of the indicated genotypes (color coding: driver control: black, responder control: grey, VPM3 neuron activation: green). Grey bars and gapped lines indicate group means ± standard deviations. *Lower panel:* Effect size analysis using estimation statistics. *Lower left side:* Mean differences (Δ) between control and test flies within each light condition (light-off: control vs test; light-on: control vs test), shown as black dots and grey bars with 95 % confidence intervals (black lines) and bootstrap distribution (grey curves). *Lower right side:* Difference of mean differences (ΔΔ), shown as black dots and grey bars with 95 % confidence intervals (black lines) and bootstrap distribution (grey curves). When ΔΔ confidence intervals exclude zero, this indicates evidence for a light-dependent effect beyond baseline differences. Genotype comparisons, light conditions, sample sizes (Ns), and effect sizes are provided in Supplementary Table 1.

### Endogenous MBON11 activity in hungry flies drives feeding in both sexes

In females, optogenetic MBON11 activation led to moderate decreases in feed count and duration, with no effects on meal size or latency—contrasting with the robust reduction in feed latency observed in males (**Figure S6E, G**). In contrast, inhibition of MBON11 in females produced very similar trends and comparable effect sizes (Δg) to males across six behavioral metrics, supporting a feeding-suppressing role in both sexes (**Figure S6F, H**). Properly controlling for genetic and light effects, an equivalent effect size (ΔΔ) in feed volume reduction (−153 nl, −120 nl) was observed in females and males, respectively (**Figure 3F, Figure S6F**). Together, these results indicate that MBON11 activation promotes feeding only in males, whereas MBON11 inhibition suppresses feeding in both sexes, indicating a sexually dimorphic aspect of this circuit.

### MBON11 synaptic output is required for octopaminergic promotion of feeding

The activation of VPMs and their postsynaptic partner MBON11 resulted in the same pro-eating trend across several Espresso metrics, leading us to hypothesize that octopaminergic feeding drive is mediated by MBON11 downstream action. Due to technical limitations (the *MB113C, MB112C*, and *XY101* drivers all use split-Gal4 combinations), we could not test MBON11 blockade in conjunction with VPM3/4 activation. As such, we used *Tdc2-LexA* to drive octopaminergic Chr expression. Consistent with the results obtained using *Tdc2-Gal4>Chr* (**Figure 1G**), *Tdc2-LexA>Chr* activation led to an increase in food consumption over 2 h, with an average +60 nl [95CI 38, 81] increase per fly (**Figure 4A**).

**Figure 4.**
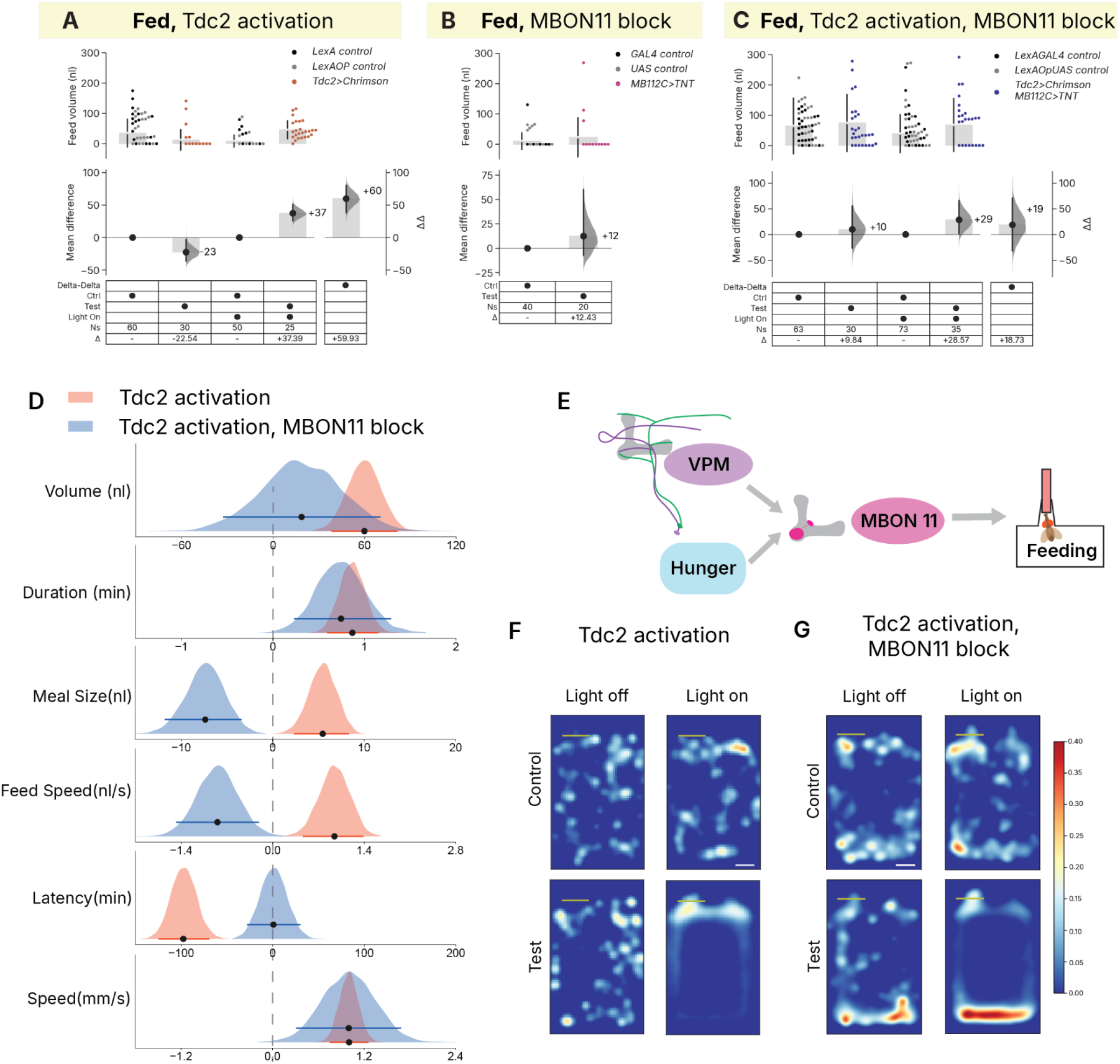
Blocking synaptic transmission from MBON11 abolishes feeding promotion induced by Tdc2 activation in several feeding and locomotion metrics. **A.** Estimation plot of feed volume for fed male flies with *Tdc2-LexA* neuronal activation. **B.** Estimation plot of feed volume for fed male flies with MBON11 synaptic-transmission block. **C.** Estimation plot of feed volume for fed male flies with *Tdc2-LexA* activation and a block of synaptic transmission in MBON11. *Upper panel:* Observations of feed volume for individual flies of the indicated genotypes (color coding: driver control: black, responder control: grey, Tdc2 neuron activation : orange red, MBON11 synaptic-transmission block: pink, *Tdc2-LexA* activation and an MBON11 synaptic-transmission block: dark blue). Grey bars and gapped lines indicate group means ± standard deviations. *Lower panel:* Effect size analysis using estimation statistics. *Lower left side:* Mean differences (Δ) between control and test flies within each light condition (light-off: control vs test; light-on: control vs test), shown as black dots and grey bars with 95 % confidence intervals (black lines) and bootstrap distribution (grey curves). *Lower right side:* Difference of mean differences (ΔΔ), shown as black dots and grey bars with 95 % confidence intervals (black lines) and bootstrap distribution (grey curves). When ΔΔ confidence intervals exclude zero, this indicates evidence for a light-dependent effect beyond baseline differences. **D.** Differences of mean differences (ΔΔ, black dots) for six different feeding-related behavior metrics for fed male flies with Tdc2-LexA activation and Tdc2-LexA activation + MBON11 synaptic block. The black bar represents the 95CI and the grey curve represents distribution. **E.** A mechanistic model for a feeding circuit involving VPMs and MBON11 neurons. **F.** Heatmaps showing the z score of time that flies (of indicated groups and conditions) spent at different locations in the *Espresso* assay fly chambers for fed male flies with *Tdc2-LexA* neuronal activation (same flies and conditions as in panel A). **G.** Heatmaps showing the z score of time that flies (of indicated groups and conditions) spent at different locations in the *Espresso* assay fly chambers for fed male flies with *Tdc2-LexA* activation and a block of synaptic transmission in MBON11 (same flies and conditions as in panel C).

MBON11 blockade alone using tetanus toxin (TeNT) did not produce any feeding-volume change in fed flies **(Figure 4B**). With the individual effects of *Tdc2-LexA>Chr* and *MBON11>TeNT* verified, we combined the two neurogenetic interventions: Tdc2-cell activation with MBON11 blockade. Compared to Tdc2 activation alone, MBON11 blockade abolished the feed-volume optogenetic effect (**Figure 4C**) and impacted several other Tdc2-induced feeding-related behaviors (**Figure 4D**). Though Tdc2-induced hyperactivity and the increase of feed duration remained, the meal-size and feed-speed effects in light-actuated *Tdc2>Chr; MBON11>TeNT* flies were reversed and the feed-latency effect was abolished (**Figure 4D**).

As observed with acute MBON11 opto-inhibition, TeNT blockade of MBON11 caused the Tdc2-activated flies to spend more time at the bottom of the chamber, even though Tdc2-cell activation on its own led to a preference for the top of the chamber near the food port (**Figure 4F, G**). The tracking data did not identify any climbing or locomotion defects in the *Tdc2>Chr; MBON11>TeNT* flies. These findings demonstrate that the behavioral effects induced by Tdc2 activation have a strong dependency on MBON11 output, supporting the hypothesis that MBON11 acts downstream of octopaminergic signaling to promote feeding (**Figure 4E**). Based on the hemibrain connectome, VPM cells are the only OA neurons that directly synapse with MBON11 (Scheffer et al. 2020). The similarity between VPM and Tdc2 activation effects on feeding metrics suggests that VPM→MBON11 signaling mediates a substantial component of octopaminergic influence on feeding behavior, though indirect pathways from other OA neurons remain possible.

### PPL101 is required for hunger-motivated feeding

As OA neurons can instruct increased feeding but are not required for normal hunger-motivated feeding behaviors, we asked whether another neuromodulatory synaptic partner of MBON11 is required for hunger-motivated feeding increases. The MB receives projections from several dopaminergic cells, with the primary source of dopaminergic input to MBON11 being the PPL101 class (**Figure 5A-D**). A connectomics analysis showed that MBON11 in the right hemisphere receives 961 synaptic connections from PPL101 in the right hemisphere and 989 synaptic connections from PPL101 in the left (**Figure 5D, Figure S9**)(Scheffer et al. 2020). These neurons have been reported to communicate satiety, and, in hungry flies, are required for persistent food-odor tracking (Krashes et al. 2009; Perisse et al. 2016; Tsao et al. 2018a; Sayin et al. 2019). Based on the prior model that PPL101 activity signals satiety, we hypothesized that PPL101 inhibition would increase feeding in fed flies while activation would reduce feeding in hungry flies. Surprisingly, both manipulations had minimal effects on food consumption (**Figure 5E, H**). In a further challenge to our hypothesis, in hungry flies, PPL101 inhibition drastically reduced food consumption for about 103 nl (−103 nl [95CI −164, −43], indicating that PPL101 activity is required for hunger-motivated feeding (**Figure 5F**). We thus hypothesized that PPL101 activity might also be instructive for feeding. Contradicting this hypothesis, however, optogenetic PPL101 activation in fed flies had little effect on feed volume or other feeding metrics (**Figure 5G, 6A**). These data indicate that PPL101 activity is required but not instructive for feeding drive, as its activity alone is insufficient to override satiety. We thus conclude that PPL101 activity has a permissive function in hunger-motivated feeding.

**Figure 5.**
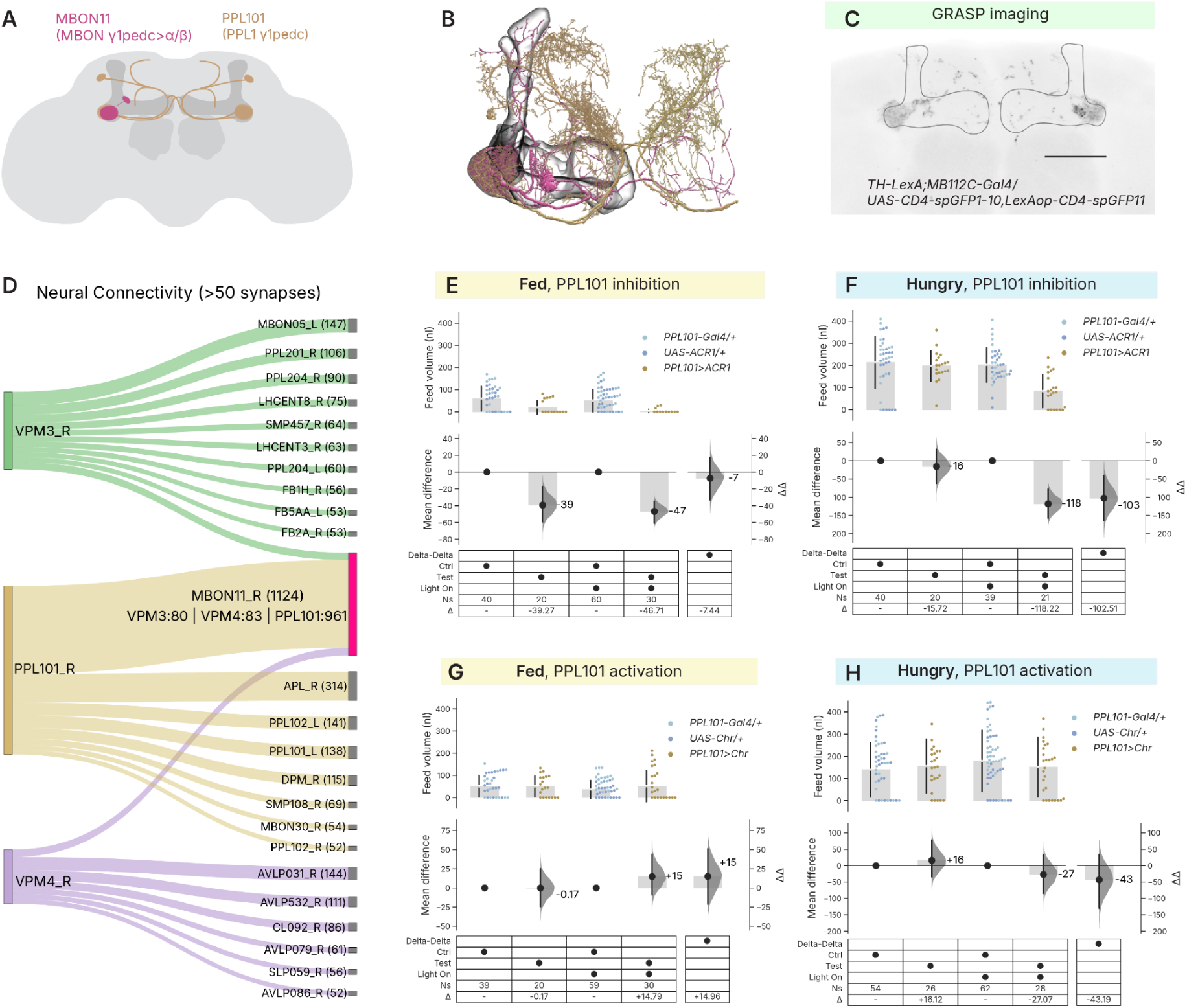
Dopaminergic PPL101 cells, presynaptic to MBON11, are required for hunger-motivated feeding. **A.** An illustration of the projection patterns of PPL101 and MBON11. **B.** A neuPrint skeleton viewer image of (Scheffer et al. 2020; Plaza et al. 2022) projections of PPL101 and MBON11 show neurite convergence in the γ1 and peduncle zones of the horizontal MB lobe. **C.** Native GRASP signal between PPL101 and MBON11 shows synaptic signal in γ1 and peduncle zones. The outlines of mushroom bodies are traced in black lines. **D.** Postsynaptic connections to VPM3, VPM4 and PPL101 (Synapse>50). Sankey plot shows the cell type of the downstream individual neurons that have strong postsynaptic connections (Synapse>50) with VPM3, VPM4 and PPL101, with the width of the link proportional to the count of the connection. The number of connections is indicated inside the brackets next to the cell types. Cell types from the left (L) or right (R) hemisphere were designated with an underscore prefix. **E.** Estimation plot of feed volume for fed male flies with PPL101 inhibition. **F.** Estimation plot of feed volume for 24-h starved male flies with PPL101 inhibition. **G.** Estimation plot of feed volume for fed male flies with PPL101 activation. **H.** Estimation plot of feed volume for 24-h starved male flies with PPL101 activation. *Upper panel:* Observations of feed volume for individual flies of the indicated genotypes (color coding: driver control: black, responder control: grey, PPL101 neuron manipulations: goldenrod). Grey bars and gapped lines indicate group means ± standard deviations. *Lower panel:* Effect size analysis using estimation statistics. *Lower left side:* Mean differences (Δ) between control and test flies within each light condition (light-off: control vs test; light-on: control vs test), shown as black dots and grey bars with 95 % confidence intervals (black lines) and bootstrap distribution (grey curves). *Lower right side:* Difference of mean differences (ΔΔ), shown as black dots and grey bars with 95 % confidence intervals (black lines) and bootstrap distribution (grey curves). When ΔΔ confidence intervals exclude zero, this indicates evidence for a light-dependent effect beyond baseline differences. Genotype comparisons, light conditions, sample sizes (Ns), and effect sizes are provided in Supplementary Table 1.

**Figure 6.**
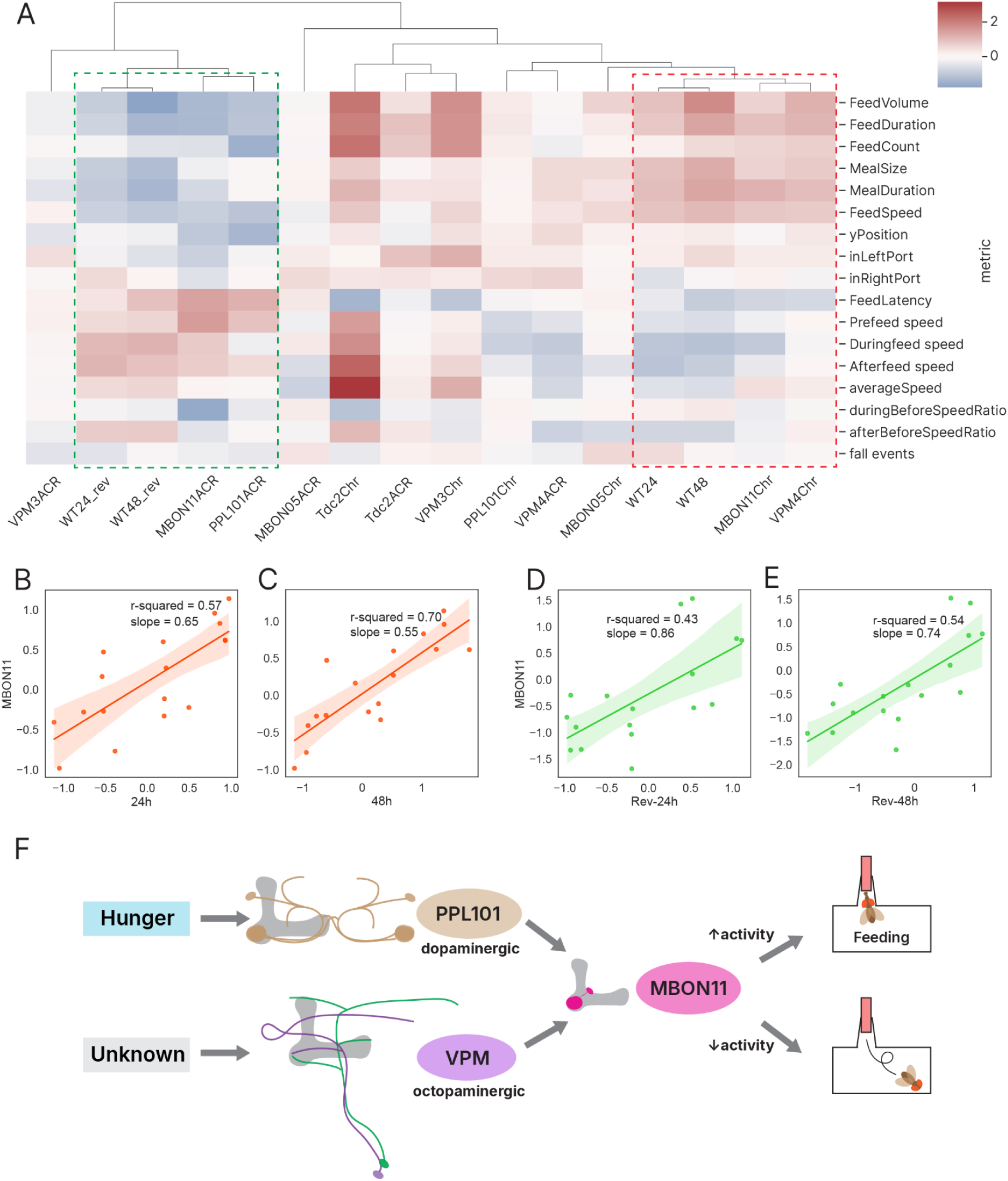
Optogenetic ethomic analysis reveals that MBON11 is essential in regulating hunger. **A.** Hierarchical clustering analysis of standardized effect sizes (Δ*g*) for 17 feeding and locomotor metrics (rows) for two starvation durations (and their inverse effects) and 12 different neuronal interventions (columns). WT24 and WT48 represent hunger-induced behavioral changes following 24-h and 48-h starvation in wild-type male flies, respectively. WT24_rev and WT48_rev represent the corresponding reversed-sign effect sizes, used to represent the hunger→satiety transition. CsChrimson activation experiments were performed in *ad libitum* fed male flies, while GtACR1 inhibition experiments were performed in 24-h starved male flies. Red dashed box: MBON11 activation and VPM4 activation cluster with WT24 and WT48. Green dashed box: MBON11 inhibition and PPL101 inhibition cluster with WT24_rev and WT48_rev. **B.** Linear-regression plot of Δ*g* of 17 feeding and locomotor metrics between 24-h starvation effects in WT flies and MBON11 activation effects in fed flies. The r value represents the Pearson correlation coefficient. Red indicates Chr activation experiments with ad libitum fed flies. **C.** Linear-regression plot of Δ*g* of 17 feeding and locomotor metrics between 48-h starvation effects in WT flies and MBON11 activation effects in fed flies. The r value represents the Pearson correlation coefficient. Red indicates Chr activation experiments with *ad libitum* fed flies. **D.** Linear-regression plot of Δ*g* of 17 feeding and locomotor metrics between reversed-sign effects of 24-h starvation in WT flies and MBON11 inhibition effects in 24-h starved flies. The r value represents the Pearson correlation coefficient. Green indicates ACR1 inhibition experiments with 24-h starved flies. **E.** Linear-regression plot of Δ*g* of 17 feeding and locomotor metrics between reversed-sign 48-h effects in WT flies and MBON11 inhibition effects in 24-h starved flies. The r value represents the Pearson correlation coefficient. Green indicates ACR1 inhibition experiments with 24-h starved flies. **F.** Mechanistic model for the feeding circuit involving VPMs, PPL101, and MBON11 neurons.

**Figure S8.**
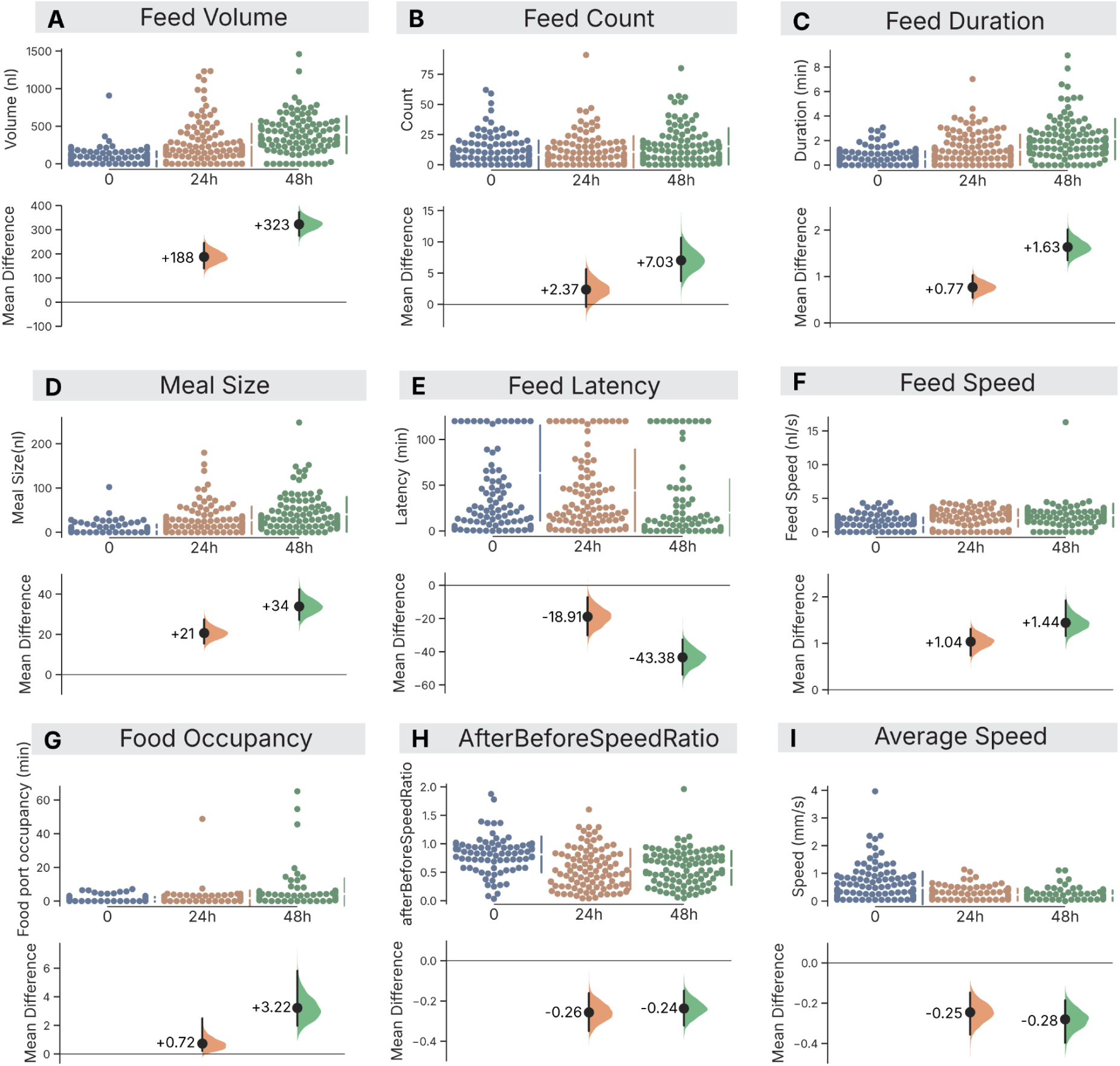
Hunger induced by 24- or 48-hs of starvation alters behavioral metrics in a 2-h Espresso assay. **A.** Estimation plot of feed volume for fed (blue), 24-h starved (orange), and 48-h starved (green) wild-type male *w^1118^*flies. **B.** Estimation plot of feed count for fed (blue), 24-h starved (orange), and 48-h starved (green) wild-type male *w^1118^*flies. **C.** Estimation plot of feed duration for fed (blue), 24-h starved (orange), and 48-h starved (green) wild-type male *w^1118^*flies. **D.** Estimation plot of meal size for fed (blue), 24-h starved (orange), and 48-h starved (green) wild-type male *w^1118^*flies. **E.** Estimation plot of feed latency for fed (blue), 24-h starved (orange), and 48-h starved (green) wild-type male *w^1118^*flies. **F.** Estimation plot of feed speed for fed (blue), 24-h starved (orange), and 48-h starved (green) wild-type male *w^1118^*flies. **G.** Estimation plot of food occupancy for fed (blue), 24-h starved (orange), and 48-h starved (green) wild-type male *w^1118^* flies. **H.** Estimation plot of peri-feed speed ratio for fed (blue), 24-h starved (orange), and 48-h starved (green) wild-type male *w^1118^* flies. **I.** Estimation plot of locomotion speed for fed (blue), 24-h starved (orange), and 48-h starved (green) wild-type male *w^1118^* flies. For each metric, the upper panel shows the observations of the respective means of individual flies (dots); the gapped lines represent the means with standard deviation. The lower panel shows the mean differences (Δ) between 24-h starvation and fed, and 48-h starvation and fed; with the 95CI (black bar) and the error distribution (colored curve).

### Hunger induces diverse changes in feeding-behavior features

To contextualize the artificial optogenetic effects on feeding, we examined how feeding behavior naturally changes in response to hunger. We subjected wild-type flies to 24-h and 48-h starvation regimes prior to Espresso recording. Data from these starved flies were compared with control, *ad libitum*-fed flies to calculate effect sizes for various feeding and locomotion metrics that result from hunger (**Figure 6A, S8**). Both 24-h and 48-h starvation increased the total food intake, meal size, feed frequency, feed duration, feed speed, and food port occupancy, while decreasing feed latency (faster feeding initiation) and reducing the walking speed (**Figure S8**). For most behavioral features, 48-h starvation produced more pronounced effects than 24-h starvation (*e.g.*, feed volume: 48h ΔΔ = +323 nl [95CI 279, 376]) vs. 24h ΔΔ= +188 nl [95CI 137, 245], indicating that the longer starvation epoch elicits a larger metabolic–behavioral state change.

**Figure S9.**
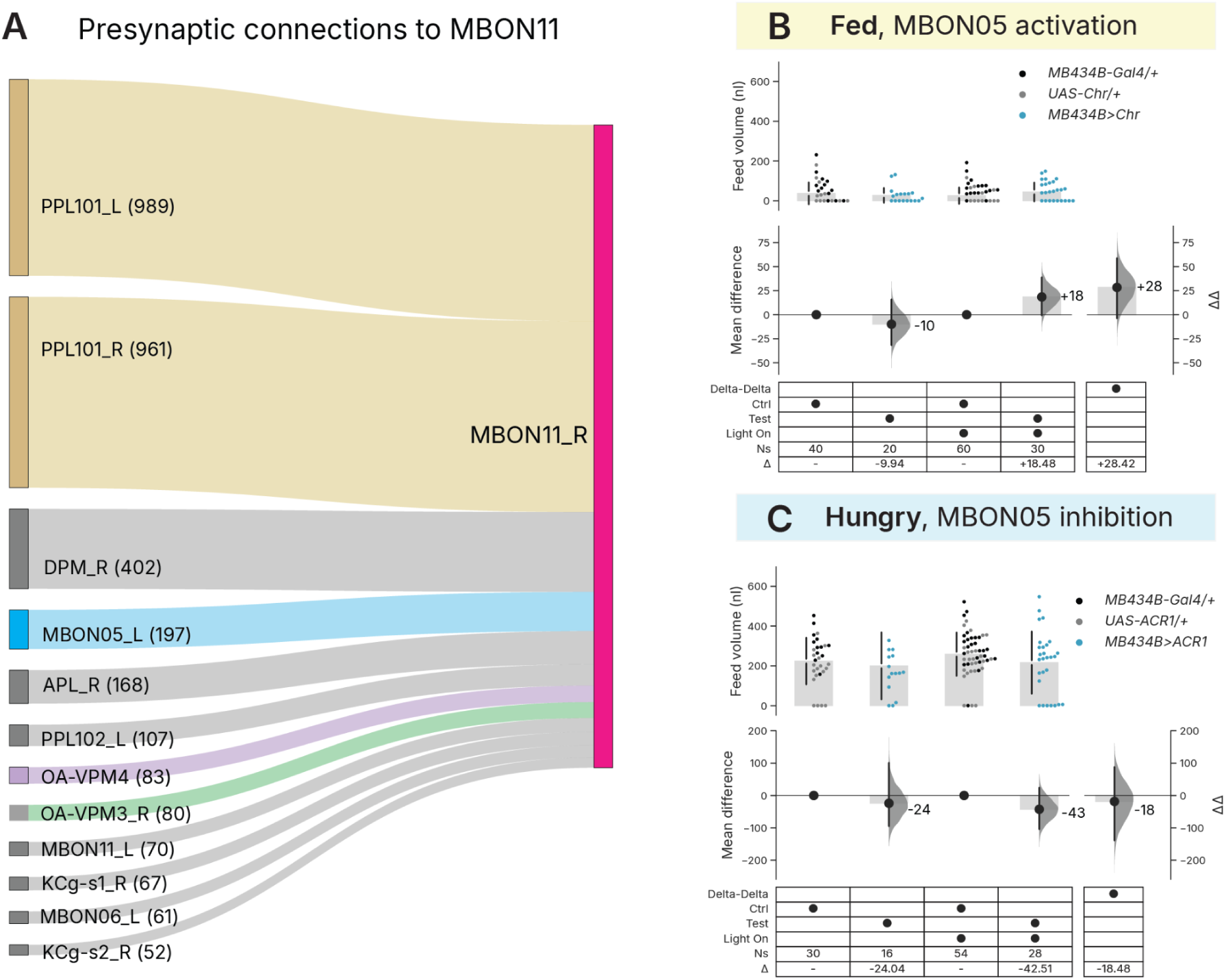
Presynaptic partners of MBON11 and MBON05 optogenetic effects on food intake. **A.** Presynaptic connections to MBON11 (Synapse>50). Sankey plot shows the cell type of the upstream individual neurons that have strong presynaptic connections (Synapse>50) with MBON11, with the width of the link proportional to the count of the connection. The number of connections is indicated inside the brackets next to the cell types. Cell types from the left (L) or right (R) hemisphere were designated with an underscore prefix. **B.** Estimation plot of feed volume for fed male flies with MBON05 activation. **C.** Estimation plot of feed volume for 24-h starved male flies with MBON05 inhibition. *Upper panel:* Observations of feed volume for individual flies of the indicated genotypes (color coding: driver control: black, responder control: grey, MBON05 neuron manipulation: blue). Grey bars and gapped lines indicate group means ± standard deviations. *Lower panel:* Effect size analysis using estimation statistics. *Lower left side:* Mean differences (Δ) between control and test flies within each light condition (light-off: control vs test; light-on: control vs test), shown as black dots and grey bars with 95 % confidence intervals (black lines) and bootstrap distribution (grey curves). *Lower right side:* Difference of mean differences (ΔΔ), shown as black dots and grey bars with 95 % confidence intervals (black lines) and bootstrap distribution (grey curves). When ΔΔ confidence intervals exclude zero, this indicates evidence for a light-dependent effect beyond baseline differences.

**Figure S10.**
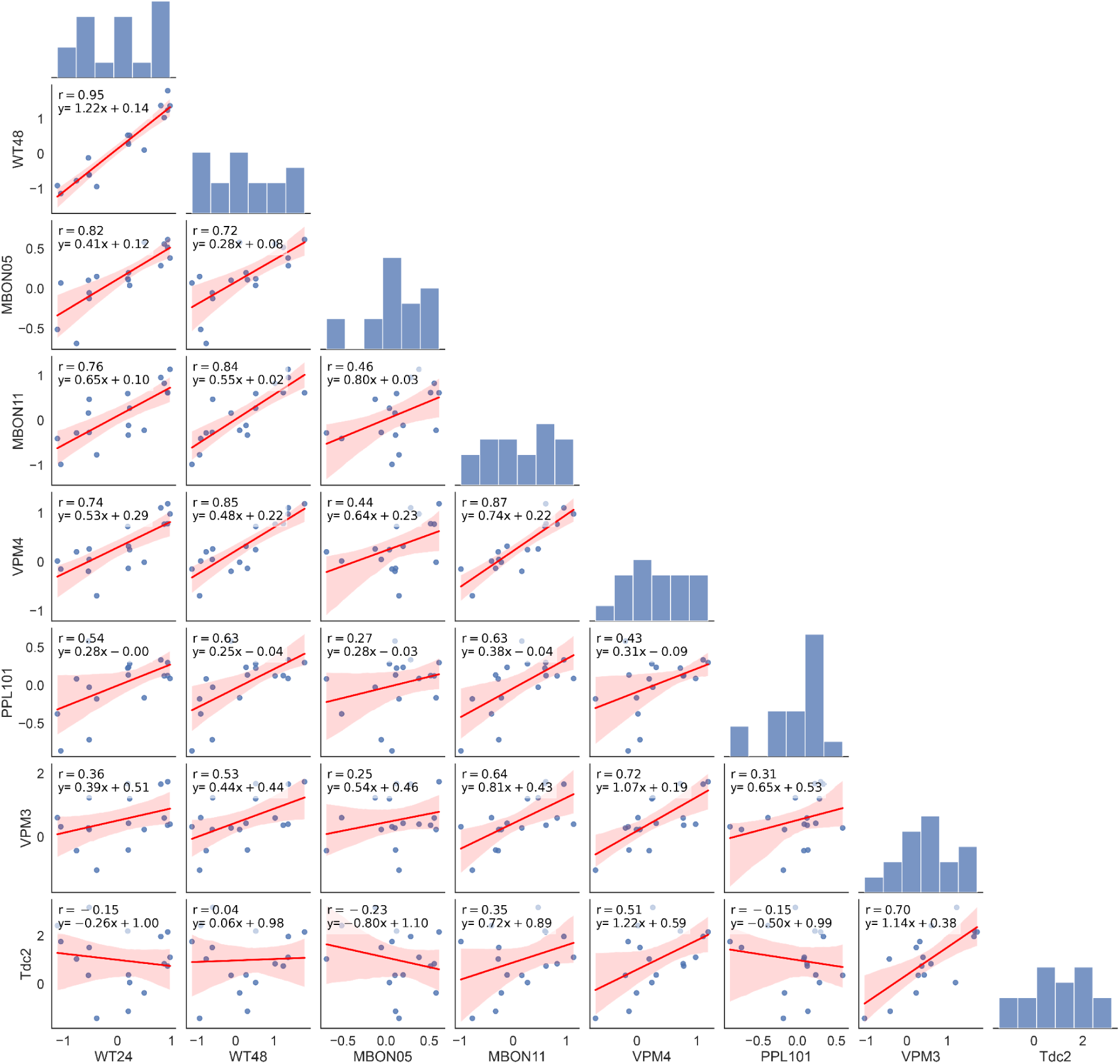
Regression pair plots of the effects of neuronal activation and starvation/satiety on behavior. Regression pair plots of Δ*g* of neuronal activation with Chr in fed flies and the wild-type starvation effect, across 17 behavioral metrics. Each dot represents the Δ*g* of a single metric for each pair of interventions. The regression line and a 95 % confidence interval for that regression are plotted in red. The histogram plots on the diagonal show the distribution of Δ*g* for each intervention. The Pearson correlation coefficient (r), and fit equation (including the slope and intercept) are printed in each panel.

**Figure S11.**
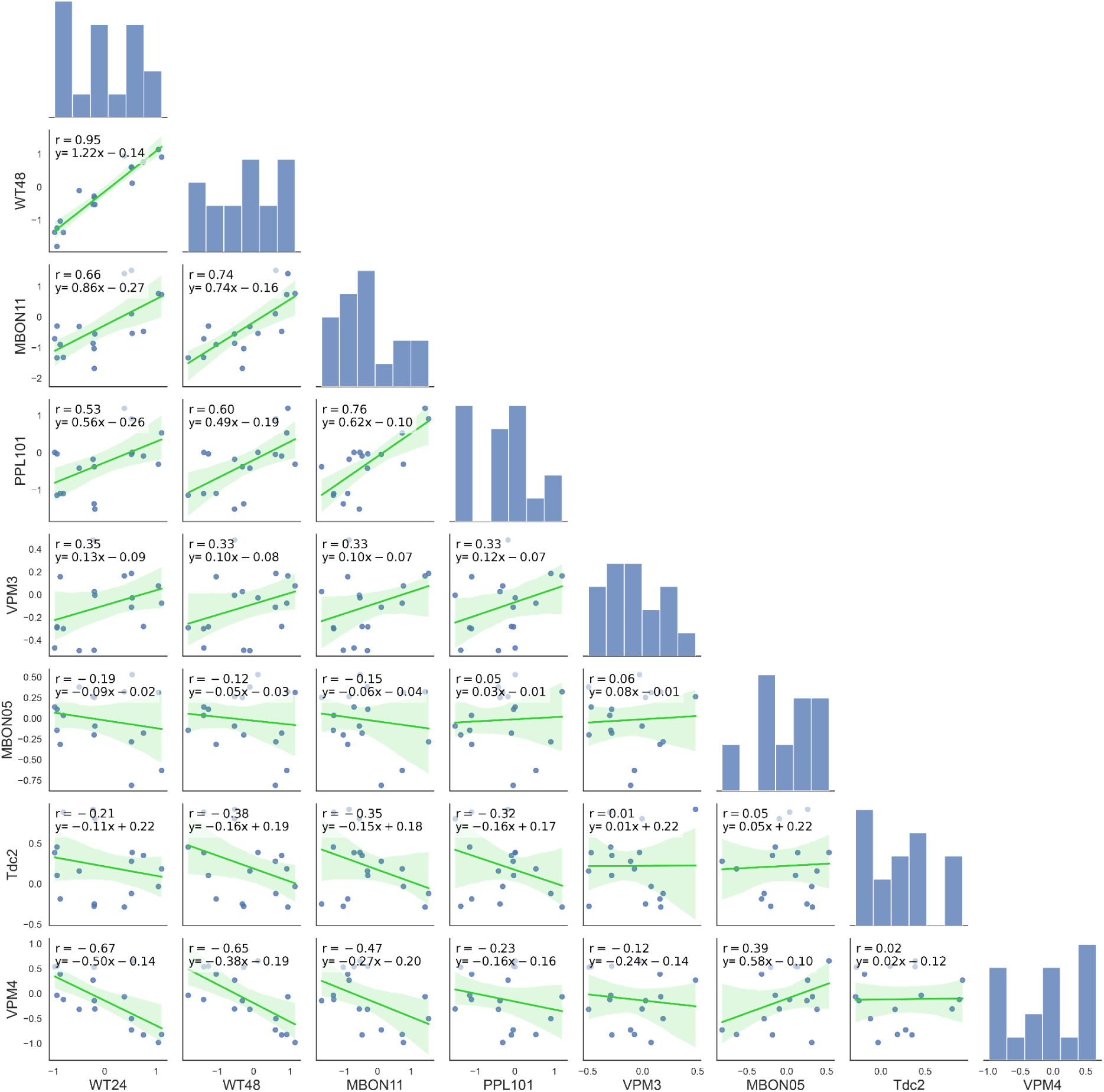
Regression pair plots of the effects of neuronal inhibition and starvation/satiety on behavior. Regression pair plots of Δg of neuronal inhibition with ACR1 in 24-h starved flies and the reverse of the wild-type starvation effect, across 17 behavioral metrics. Each dot represents the Δ*g* of a single metric for each pair of interventions. The regression line and a 95 % confidence interval for that regression are plotted in green. The histogram plots on the diagonal show the distribution of Δ*g* for each intervention. The Pearson correlation coefficient (r), and fit equation (including the slope and intercept) are printed in each panel.

### Effect-size clustering shows that most optogenetic interventions are distinct from hunger

Our preceding analysis focused on eating as the key indicator of feeding behavior; however, we also considered that food consumption could be altered by possible optogenetic pleiotropic effects in circuits that normally play only a minor role in the natural hunger state. Given that overall feeding behavior encompasses multiple features, we adopted an ethomics approach to analyze the phenotypic effect-size vectors (‘phenovectors’) of each intervention (Robie et al. 2017). We hypothesized that circuits genuinely involved in feeding control would be similar to natural hunger–satiety effect vectors, while circuits with incidental effects on feeding would not. To generate effect-size vectors with diverse metrics, we transformed each into a Hedges’ *g* value, a standardized measure of effect size (Borenstein et al. 2009). Each experiment produced a vector of 17 *g* values for food volume, walking speed, and 15 other metrics. Hierarchical clustering was conducted on 12 optogenetic experiments and two starvation experiments. The starvation effects were used in two ways: representing both the sated→hungry and, by sign reversal, the hungry→sated transitions. Clustering placed the 24-h and 48-h hungry↔sated profiles together (**Figure 6A**); this similarity was supported by linear regression, which revealed a strong correlation between the 24- and 48-h hungry↔sated vectors (*r* = 0.95; **Figure S10, S11**).

While the clustering analysis showed that the natural hunger (and satiety) vectors grouped together, most of the optogenetic interventions formed distinct outgroups (**Figure 6A**). This was even true for lines that elicited robust feeding increases, including *VPM3>Chr* and *Tdc2>Chr*. These two lines showed feeding-metric changes that in some cases, exceeded those observed after 48-h starvation, while some of their speed effects were opposite to those seen in hunger profile. Most of the other neuromodulatory experiments were dissimilar to the hungry/sated profiles (**Figure 6A**).

To contextualize the optogenetic phenotype vectors, we also tested MBON05 neurons (also known as MBON-γ4>γ1γ2) and included it in the clustering analysis. MBON05 is a postsynaptic partner of VPM3 (147 connections) and also a presynaptic partner of MBON11 (197 connections) (**Figure 5D, Figure S9A**) (Scheffer et al. 2020; Plaza et al. 2022). MBON05 interventions did not affect food consumption (**Figure S9B, C**). Although the MBON05*>Chr* phenotypic vector clustered in proximity to the sated→hungry vector, *MBON05>ACR1* phenotypic vector failed to cluster with the hungry→sated vector (**Figure 6A**).

These findings suggest that while optogenetic activations of Tdc2, VPM3 and VPM4 cells all robustly increase food consumption, the overall behavioral profiles differ markedly from those associated with natural hunger states. These findings are consistent with the idea that these neurons promote feeding by eliciting internal states that are dissimilar to authentic, starvation-induced hunger.

### Optogenetic ethomics indicates that MBON11 controls the hunger–satiety state change

The cell type that most closely phenocopied natural hunger profiles was MBON11 (**Figure 6A-E**). Activation of MBON11 using *MBON11>Chr* clustered next to the sated→hungry effects (**Figure 6A**). Similarly, MBON11 activation closely correlated with the hunger vectors, with *R^2^* values of 0.57 and 0.70 for 24- and 48-h starvation, respectively (**Figure 6B, C**). Moreover, inhibition with *MBON11>ACR1* clustered next to hungry→sated effects (**Figure 6A**), while MBON11 inhibition correlated with the hungry→sated effects, with *R^2^* values of 0.43 and 0.54 for 24- and 48-h ‘reverse starvation’, respectively (**Figure 6D, E**). We also examined regression slopes, where a slope of 1.0 would indicate a perfectly direct relationship. For activation, the MBON11∼starvation slopes were moderate (0.65, 0.55, **Figure 6B, C**), while for inhibition, the MBON11∼satiation slopes were relatively stronger (0.86, 0.74, **Figure 6D, E**). These observations are consistent with the idea that MBON11-neuron activity exerts control over the hungry↔sated state transitions. The *R^2^* and slope estimates indicate that MBON11 is not solely responsible for these transitions; nevertheless it can account for around 50 % of behavioral changes along the hungry↔sated axis. Given that MBON11 activation further enhances feeding in 24-h starved flies (**Figure S6C**), and that the MBON11-activation effect profile shows a better correlation with the 48-h hunger vector (*R^2^* = 70%, **Figure 6B, C**), we infer that MBON11 activation induces a hunger-like state more intense than that produced by 24-h starvation.

The other two optogenetic lines that clustered close to the hunger/satiation profiles were *VPM4>Chr*, which was similar to the sated→hungry transitions, and *PPL101>ACR1*, which grouped with the hungry→sated transitions (**Figure 6A**). These lines were also similar to the MBON11 phenovectors, with *R^2^* values of 0.76 and 0.58, respectively (**Figure S10, S11**). These observations lend further support to the above-mentioned hypotheses that VPM4 activity can instruct a hunger-like state, while PPL101 quiescence can induce a satiety-like state.

## Discussion

This study reveals a neural circuit connecting through the *Drosophila* mushroom body that integrates octopaminergic and dopaminergic signals to control feeding behavior. With optogenetic manipulations and systematic behavioral analyses, we demonstrate that four interconnected cell types—VPM3, VPM4, PPL101, and MBON11—converge at a critical circuit node to regulate food consumption. Our findings establish MBON11 as a bidirectional regulator of feeding that receives inputs from octopaminergic VPM neurons, which promote feeding when activated but are dispensable for hunger-driven consumption, and dopaminergic PPL101 neurons, which are essential for hunger-motivated feeding but cannot independently override satiety. Quantitative ethomic analysis reveals that MBON11 activity can account for approximately half of the variance of the hunger–satiety state vector, positioning MBON11 as an integrator of diverse neuromodulatory signals into a coherent feeding state. These results refine views of octopamine’s role in hunger and reveal how mushroom body circuits, classically associated with learning and memory, directly control consummatory behaviors through state-dependent neuromodulation.

### VPM4 activity promotes feeding initiation in females

OA has long been proposed to promote feeding and feeding-related behaviors (Zhang et al. 2013; Yang et al. 2015; Li et al. 2016; Tian and Wang 2018; Youn et al. 2018). Of the two VPMs, feeding research has most studied the OA-VPM4 cells. In females tested for taste-induced proboscis extension, VPM4 activity increased proboscis-extension probability, interpreted as promoting feeding initiation (Youn et al. 2018). Consistent with this, our experiments show that activating VPM4 in fed flies reduces feeding latency and increased feed count (**Figure S2H**), i.e. accelerated feeding initiation. The effects of VPM4 inhibition reveal sex differences: in females, inhibition impeded feeding initiation (decreased feed count) while also increasing meal size and leaving total consumption unchanged (**Figure S3H**); males showed minimal response to VPM4 inhibition, with a moderate increase in meal size but no effect on feed count (**Figure S3H**). The two sets of findings from proboscis extension (Youn et al. 2018) and Espresso (this study) are concordant in their support of VPM4’s role in female feeding initiation. However, since consumption is unaffected in light-actuated VPM>ACR1 flies, our data demonstrate that an individual feature (*e.g.* initiation) cannot be used as a proxy for the overall behavioral program.

### VPM4 effects on locomotion depend on behavioral paradigm

Previous work with a ball-walking assay showed that activation of Tdc2 or VPM4 neurons in hungry flies reduced vinegar-odor-evoked locomotion, interpreted as an OA brake on food-odor tracking (Sayin et al. 2019). By contrast, in freely moving flies in Espresso, Tdc2 activation drove a substantial increase in locomotion, including pre-feed locomotion (**Figure 1H, S1F, 6A)**, whereas VPM4 activation drove increased overall locomotion in starved male flies and fed female flies (**Figure S2D, H, Figure 6A**). Rather than a brake on feeding-related behaviors, octopaminergic activation led to flies locating food faster, larger meal sizes, and more total consumption (**Figure 6A)**. These findings indicate that VPM4 promotes feeding and foraging in freely moving flies, though reconciling and generalizing results across different behavioral paradigms will require further investigation.

### VPM3 exhibits a distinct feeding-regulatory profile

VPM3 activation modulates feeding behavior and promotes consumption directly, and has notable differences from VPM4. The two neuronal types showed different optogenetic phenotype profiles in the clustering and regression analyses (**Figure 6A, S10**). In the clustermap (**Figure 6A**), the VPM3 activation profile aligns more closely with broad-Tdc2-neuron effects, while VPM4 activation is similar to MBON11 activation. This analysis thus indicates that VPM3 and VPM4, despite their anatomical similarity, produce distinguishable patterns of effects on feeding and locomotor behaviors.

### Broad OA activity has smaller effects than VPM activation

The similar magnitude of feeding-related effect sizes between Tdc2 activation and VPM activations suggests that VPM3 and VPM4 may be the major OA feeding neurons. However, there are important differences: unlike the VPMs, the Tdc2 driver can induce large increases in locomotion speed, possibly mediated by non-VPM cells. Tdc2 activation increased feed volume by +68 nl [95CI 49, 90], while VPM3 activation increased it by +102 nl [95CI 63, 140] and VPM4 by +80 nl [95CI 38, 121]. VPM3 and VPM4 activations thus produced feed volume increases similar to or exceeding those from broad Tdc2 activation, which we interpret to reflect OA-cell heterogeneity. Previous work has shown that distinct OA neurons can have opposing effects on behavior (Zhang et al. 2013; Claßen and Scholz 2018), so simultaneous activation of all OA neurons may engage competing circuits or lead to downstream receptor saturation, limiting the net Tdc2 effect.

### Octopaminergic activity is not required for hunger-motivated feeding

Although activation of octopaminergic neurons (Tdc2 pan-OA, VPM3, or VPM4) in fed flies elicited substantial increases in feeding, inhibition of these neurons in starved flies did not diminish food intake. This indicates that OA activity is not acutely required for post-fasting, hunger-motivated feeding. Previous evidence that *Tbh^nM18^* mutant flies exhibited reduced sucrose intake (Li et al. 2016; Berger et al. 2024) may represent a chronic trait distinct from the hunger–satiety cycle, perhaps related to reduced locomotion and lower energy consumption. VPM inhibition did affect locomotion in hungry flies; VPM4 inhibition led to a reduction in the average locomotion speed in both sexes whereas VPM3 inhibition had no strong effect on locomotion speed (**Figure S3G-H**). Although VPM3/4 do not appear to encode hunger or satiety, these neurons likely interact with nutritional state and insulin signaling pathways.

### Potential mechanisms underlying octopaminergic feeding effects

The context in which VPM activity influences feeding behavior remains unclear. One possibility is that OA-neuron activity drives a non-hunger program that has secondary effects on feeding. First, VPM likely modulates responses to sensory stimuli. OA has been shown to promote stimulus dishabituation in both locusts and flies (Sombati and Hoyle 1984; Bacon et al. 1995; Suver et al. 2012; Scheiner et al. 2014). One group has demonstrated that VPM4 activity potentiates sugar sensing (Youn et al. 2018), while another has shown that VPM3 responds to both sugar ingestion and odor presentation in calcium imaging experiments, with these responses declining upon repeated exposure (Kapoor and Waddell 2024). Thus, VPM activation might enhance the salience of food taste and odor cues, causing dishabituation and prompting sated flies to feed repeatedly.

Another possibility is that VPM may function as a context-dependent feeding switch. Octopamine has been implicated in action selection: previous studies have demonstrated that distinct octopaminergic cell types can shift olfactory valence from neutral to either attractive or aversive (Claßen and Scholz 2018), and modify visual object preferences from avoidance to approach (Cheng et al. 2019). In male-specific behaviors, octopamine shapes behavioral hierarchies, promoting aggression over courtship (Certel et al. 2007) and courtship over sleep(Machado et al. 2017). Consistent with these findings, VPM activity could modulate arousal and prioritize feeding in the context of food availability; specifically, VPM4 might regulate motor preparation for feeding behavior.

### Dopaminergic PPL101 activity has a permissive role in hunger-driven feeding

The PPL1 dopaminergic neurons have been investigated for their roles in aversive olfactory memory (Schwaerzel et al. 2003; Riemensperger et al. 2005; Claridge-Chang et al. 2009; Aso et al. 2012), and feeding-related behaviors (Krashes et al. 2009; Aso et al. 2012; Tsao et al. 2018a; Sayin et al. 2019). The PPL101 (PPL1-γ1pedc, MB-MP1) neurons are critical for associating electric shocks with odors, with activity that correlates with shock intensity and uniquely responding to shock-conditioned odors (Vrontou et al. 2021; Villar et al. 2022). The PPL101 cells play a role in postprandial memory consolidation (Musso et al. 2015; Pavlowsky et al. 2018). Other findings indicate that PPL101 activity increases energy metabolism in MB after spaced training (Plaçais et al. 2017), that its activity changes with metabolic state (Meschi et al. 2024), and that it influences food-odor-evoked locomotion (Sayin et al. 2019).

In learned odor avoidance, PPL101 activity is both necessary and sufficient to induce learning (Aso et al. 2012). In our optogenetic inhibition experiment, silencing PPL101 neurons in hungry flies decreased consumption, acutely disrupted hunger-motivated behaviors, and broadly phenocopied the hunger→satiety transition, suggesting that hungry-state PPL101 activity supports food-seeking and consumption. However, in fed flies, optogenetic activation of these neurons did not increase feeding. From these results we infer that PPL101 neurons are active during hunger and are required to maintain that state. Thus, PPL101 activity appears to serve a permissive function for feeding: it is required for a large fraction of hunger-motivated consumption, but artificially induced activity is unable to override satiety.

### Reconciling PPL101 feeding effects across studies

Our PPL101 findings speak to prior work on these cells and feeding-related behaviors (Supplementary Table 2) (Krashes et al. 2009; Tsao et al. 2018a; Sayin et al. 2019; Meschi et al. 2024). Consistent with all previous studies, we find that PPL101 functions depend on metabolic state and that the PPL101→MBON11 connection is relevant to feeding. Our observation that PPL101 inhibition in hungry flies reduces consumption aligns with the prior finding that PPL101 inhibition suppresses persistent food-odor tracking (Sayin et al. 2019). However, prolonged optogenetic activation of PPL101 had no effect on consumption in either nutritional state, contrasting with reports that brief thermogenetic activation of PPL1 cells in starved flies suppresses appetitive memory (Krashes et al. 2009) and reduces food-seeking latency (Tsao et al. 2018a). Such apparent contradictions may arise from different behavioral paradigms and readouts, or differences between optogenetic and thermogenetic interventions. Calcium-imaging methods showed that starvation suppresses baseline PPL101 activity, though the cells maintain robust phasic responses to shock during aversive reinforcement (Meschi et al. 2024), suggesting that PPL101’s natural functions may require precisely timed activity patterns that continuous optogenetic stimulation cannot replicate. Our findings position PPL101 as providing permissive, state-dependent gating of hunger-motivated behaviors. Nevertheless, there remain outstanding questions about possible differences between learned and innate behaviors, olfactory attraction versus food consumption, and diverse methodologies that fall beyond the scope of the present study (Supplementary Table 2).

### The role of MBON11 as a mediator of the hunger behavior pattern

Previous studies have implicated MBON11 activity in various behaviors, primarily innate and learned olfactory behaviors. In optogenetic self-administration assays, MBON11 activity was found to be acutely attractive (Aso, Sitaraman, et al. 2014; Mohammad et al. 2022). Another study showed that activating MBON11 facilitated the formation of appetitive memory for an odor, while inhibiting MBON11 produced a punishing effect (König et al. 2019). MBON11 activity is required for the expression of short-term memories (Perisse et al. 2016), and silencing MBON11 during unpaired-odor training abolished memory formation (Ueoka et al. 2017). In hungry flies, MBON11 neurons display increased odor-evoked activity (Perisse et al. 2016; Tsao et al. 2018a). Furthermore, two studies found that inhibiting MBON11 synaptic output in hungry flies reduces food-odor-evoked locomotion (Tsao et al. 2018a; Sayin et al. 2019), while Sayin *et alia* showed that MBON11 activation in fed flies increases odor-evoked locomotion (Sayin et al. 2019). These diverse findings indicate that MBON11 participates in linking olfactory signals with motivational states, including hunger.

In this study, we demonstrate for the first time that MBON11 activity and quiescence directly affect feeding consumption. A diminished odor cue did not weaken this influence, suggesting that the effects of MBON11 on odor-evoked behaviors may derive from its role in feeding control. Moreover, induced MBON11 quiescence elicited an overall behavioral profile similar to the hunger→satiety transition itself, further supporting the idea that MBON11 drives the hunger behavioral state. As optogenetic interventions do not fully phenocopy the natural energy axis, it is likely that other factors are required to coordinate hunger↔satiety behavioral transitions, possibly including other MBONs, neurons in other parts of the nervous system, and/or hormones. While MBON11 activation promotes feeding only in males, MBON11 inhibition under hunger conditions in both sexes reduces feeding to levels similar to that in sated flies. These findings position MBON11 as a critical effector of hunger-driven feeding.

### The relationship between the neuromodulatory and MBON11 neurons

The feeding effects of Tdc2-cell activation were dramatically altered by blocking MBON11 synaptic transmission, demonstrating that octopaminergic feeding promotion requires MBON11. Their functional interaction, however, is not necessarily the result of direct synaptic connections. While VPM3/4 cells are the only OA neurons known to directly synapse with MBON11 neurons in the γ1 and peduncle zones, they also have arborizations that extend broadly, even within the MB (Busch et al. 2009). Thus, there could be several layers of intermediary synaptic connections and therefore indirect pathways might contribute to this functional circuit. Additionally, the hemibrain connectome represents a snapshot of a single adult brain and may not capture all anatomical connections or variability across individuals (Scheffer et al. 2020). Furthermore, state-dependent or modulatory connections that depend on neuromodulatory context or physiological state may not be revealed by structural connectomics alone (Benefiel and Greenough PhD 1998; Kremer et al. 2010; Nelson et al. 2024). While connectomic, GRASP, and epistasis data strongly support direct VPM→MBON11 connectivity mediating feeding effects, electrophysiological recordings of synaptic transmission would lend further, functional support to these conclusions.

Reconciling the various roles of the PPL101→MBON11←VPM nexus in aversive and appetitive conditioning, memory consolidation, and feeding control might lie in its overarching function in managing energy stress. Both appetitive-learning protocols and most post-starvation feeding-related behavior protocols employ extended periods of starvation (14 h or more), which induces energy stress (Krashes and Waddell 2008; Tsao et al. 2018a; Sayin et al. 2019). Odor-shock conditioning displays the molecular hallmarks of energy stress (Plaçais et al. 2017). Shock stress, like hunger, may induce energy stress by a combination of increased locomotor activity, and/or energy expenditure to restore membrane gradients in the nervous system after systemic depolarization. We speculate that the MBON11 nexus has an essential role in responding to energy stress, by promoting behaviors that restore energy homeostasis, both in the contexts of acute hunger and olfactory-memory acquisition to guide future foraging.

### Study limitations

Several limitations should be considered when interpreting our results. Although we controlled for motor function, the vertical assay design could misinterpret climbing deficits as reduced feeding motivation. The 2-h assay window cannot capture chronic internal states that might persist for days. Our continuous light actuation may not accurately recapitulate the natural firing pattern of neurons; *e.g.* PPL101 might require specific activity patterns that continuous activation cannot replicate. As with all optogenetic manipulations, artificial activation could produce effects that differ from natural physiology. The enhanced feeding from OA activation, for example, could be neomorphic, as the natural contexts that activate VPM neurons to drive increased feeding remain unknown. We are unable to completely explain why the Tdc2-activation effects are not the sum of the two VPM cell effects; this question could eventually be addressed ‘by silencing VPMs with TeNT under the control of the VPM3 and 4 drivers’ during Tdc2-LexA activation. We are also unable to explain why different studies observe differences in the effects of PPL101 on feeding-related behaviors. Future studies using further epistasis manipulations, chronic tracking, additional action-sequence analyses, physiological recordings in behaving flies, additional behavioral paradigms, and/or temporally patterned stimulation could address these limitations.

### Outlook

Using direct feeding measurement, phenovector analysis, and state-contextualized optogenetic ethomics, we demonstrate that four connected cell types all control feeding in specific ways: VPM3 and VPM4 can drive feeding in sated flies but aren’t needed for hunger-driven consumption; PPL101 enables hunger-driven feeding; and MBON11 can instruct food consumption in both directions (**Figure 6F**). MBON11 is required for octopaminergic feeding effects, and its activity and quiescence phenovectors match those of natural hunger-satiety transitions.

The PPL101→MBON11←VPM circuit motif illustrates how innate and learning elements are fundamentally intertwined with feeding control. Recent studies and our evidence support that the mushroom body serves as a central hub where sensory, motivational, and reinforcement signals are integrated to guide feeding behaviors (Tsao et al. 2018b; Suárez-Grimalt et al. 2024). Within this architecture, MBON11 appears to function as a convergence point linking memory and motivational signals to modulate consumption. Comparable architectures are observed in mammals, where hypothalamic circuits integrate cortical, limbic, and neuromodulatory inputs to coordinate feeding and learning (Sternson 2013). These cross-species parallels suggest conserved computational strategies for flexible, context-dependent regulation of food intake.

## Materials and Methods

### Drosophila husbandry

Flies were reared on standard cornmeal-based food medium at 25°C, and 12-h light–12-h dark cycle, except for flies used in the optogenetic experiments. For optogenetic experiments, flies were kept in the dark in vials or bottles wrapped in aluminum foil throughout their development. These flies were raised on standard food and transferred onto 0.5 mM all-trans-retinal (ATR; Sigma-Aldrich, USA)-supplemented food 2 days prior to the experiments.

### Fly stocks

Wild-type flies were from a cantonized *w^1118^* line. The *Tdc2-lexA:VP16* line was kindly provided by Dr. Sarah J Certel (Andrews et al. 2014). The OA split-Gal4 line *MB113C-Gal4* (VPM4 neurons), the MBON05 line *MB434B-Gal4*, the PPL101 line *MB320C-Gal4* and the MBON11 line *MB112C-Gal4* were kindly provided by Gerry Rubin (Janelia Research Campus) (Aso, Hattori, et al. 2014; Aso, Sitaraman, et al. 2014). The following lines were obtained from the Bloomington Drosophila Stock Center (BDSC): *Tdc2-Gal4* (RRID:BDSC_9313), *Tbh^nM18^* (RRID:BDSC_ 93999) (Monastirioti et al. 1996), *20x-UAS-CsChr* (RRID:BDSC_55135) (Klapoetke et al. 2014), *20x-UAS-CsChr* (RRID:BDSC_55136), *13XLexAop2-IVS-CsChrimson.mVenus* (RRID:BDSC_55139), *Tdc2-p65.AD* (RRID:BDSC_70862) (FlyBase 2018), *R76H04-Gal4.DBD* (RRID:BDSC_75633) (Dionne et al. 2018), *UAS-CD4-spGFP1–10, lexAop-CD4-spGFP11* (GRASP) (RRID:BDSC_ 58755) (Gordon and Scott 2009) and *UAS-TeNT* (RRID:BDSC_28837) (Sweeney et al. 1995). The *UAS-GtACR1* line was generated previously in our lab (Mohammad et al. 2017). The following lines were obtained from the Vienna Drosophila Resource Center (VDRC): *UAS-vGlut1-RNAi* (VDRC 104324), *UAS-Tbh-RNAi* (VDRC 107070) (Dietzl et al. 2007). Specific labeling of VPM3 neurons was achieved by generating a new split-Gal4 line using *Tdc2-p65.AD (FlyBase 2018)* and *R76H04-Gal4.DBD* (Dionne et al. 2018), termed XY101.

### Preparation of transgenic animals

The test flies that were carrying both driver and responder genes were compared with two otherwise isogenic control fly lines, each carrying only one of the transgenes. Control animals were generated by crossing the driver or responder line with wild-type *w^1118^* flies. All three optogenetic crosses were kept at 25°C and covered with aluminum foil to block light and prevent actuation of the opsin. Progeny F1 male or mated female flies were maintained in groups of 15–20 in vials for 5–10 days before performing behavioral assays. These fly vials were foil-wrapped and kept at 25°C until the assay.

### Feeding experiments with Espresso

An automated capillary-based custom feeding experiment system called Espresso was used to collect feeding data (Ott et al. 2024). The design for the Espresso chips was based on the CAFE assay (Ja et al. 2007; Murphy et al. 2017; Yapici et al. 2016). In brief, individual flies were housed in custom-fabricated plastic chips, each containing five feeding chambers. A cover slip was stuck to the chip by magnets at the two ends. Flies in each chamber were provided with food through two feeding tube channels, each containing a glass capillary. Experiments were performed using a panel assembly holding six Espresso chips. The Espresso panel was put in an incubator in the upright position during the behavior assay. A camera was placed in front of the platform to record fly behavior in all the chips. A video of the entire panel was captured at a rate of 30 fps by a camera assembly comprising a 4.4–11 mm high-resolution varifocal lens, a CCD camera, and cabling that transmitted data to an acquisition computer. With the CRITTA LabVIEW software, the Espresso system was able to track the locomotion of the flies, the start and end of each feeding event, and the amount of food consumed in each feeding event. In this study, fed (or ‘sated’) flies were provided food *ad libitum*, while hungry flies were starved for 24 h prior to feeding assays. Note that *ad libitum*-fed flies will have some small variation in their satiation depending on where they are in the feeding (or snacking) cycle.

### Optogenetic experiments

Optogenetic feeding experiments were performed as previously described (Ott et al. 2024). Briefly, Chr or GtACR1 was used to activate or inhibit (respectively) the targeted neurons, respectively, and the flies were illuminated using red (peak emission 617 nm, Luxeon Rebel, SR-05-H2060, Quadica Developments Inc.) or green (peak emission 530 nm, Luxeon Rebel, SP-05-G4, Quadica Developments Inc.) LEDs during feeding sessions to control neuronal activity. Experiments in the absence of neuronal activation were conducted in the dark.

Infrared back lights were used to illuminate the Espresso chips for tracking. The liquid food consisted of 5 % w/v sucrose and 5 % w/v yeast extract (Bacto Yeast Extract, Gibco #212750). To mark the food level under infrared light, a small amount of meniscus marking dye (Murphy et al. 2017) was siphoned into the capillary above the liquid food, and the position of the interface between the marking dye and liquid food was used to track changes in the food level. Male flies were collected and maintained on normal food for 3 days, before being transferred onto food containing 0.5 mM ATR for 2 days. For feeding experiments with hungry flies, the flies were starved for 24 h on 2 % w/v agarose before the behavior assays. The flies were then transferred to the Espresso apparatus, and the experiment was conducted for 2 h.

### Ground feeding assay

Male flies aged 5–10 days old were starved for 24 h before being transferred into vials with sucrose and food-color-soaked (Bake King Liquid Food Colours, Cherry Red) Kimwipe tissues embedded in the agarose floor. The vials were placed in an incubator at 25°C and exposed to a constant LED green light for 2 h. The number of flies with red-dyed (Tanimura et al. 1982) sucrose liquid in their abdomen after 2 h of feeding was visually determined using a stereomicroscope.

### Espresso: feeding event detection and measurement

The system tracked liquid food levels by imaging an infrared (IR)-absorbing dye band marking the meniscus in each capillary (Murphy et al. 2017). Simultaneously, individual fly positions were tracked via centroid detection at 30 frames per second (CRITTA software written in LabVIEW). Feeding events were identified when two criteria were met: (1) fly presence within a virtual feed port (defined as a 2 mm region from the capillary tip), and (2) food level reduction exceeding the baseline evaporation rate. Feed events were terminated when either the food level stabilized or the fly exited the feed port. To distinguish genuine consumption from noise, food level changes were processed using a moving average convergence/divergence (MACD) algorithm. Real-time evaporation rates were continuously measured from unoccupied capillaries and subtracted from detected feed volumes.

### Espresso data acquisition

The CRITTA video-tracking software Espresso plugin (LabVIEW) simultaneously monitored up to 30 individual flies across separate chambers, each containing one capillary and one fly. For each detected feed event, the system recorded: timestamp, consumed volume, event duration, concurrent evaporation rate, fly position coordinates (X,Y), and instantaneous velocity. All behavioral data from CRITTA were further processed using custom Python software (esploco, available at https://github.com/sangyu/esploco).

### Feeding analysis and ethomics

The Hedges’ *g* (Hedges 1981) of effect sizes for 17 behavioral metrics computed by the Espresso feeding and locomotion analysis package ESPRESSO (https://github.com/ACCLAB/espresso) and Esploco (https://github.com/sangyu/esploco) were built using the Dabest Estimation statistics python package (https://github.com/ACCLAB/DABEST-python) (Ho et al. 2019). Hierarchical clustering analysis was performed using the Seaborn clustermap function using the “correlation” metric (Waskom 2021).

### Espresso behavioral metrics

The following feeding metrics were quantified:

- **Feed volume**: Total volume of liquid food consumed during the assay period (nL)
- **Feed count**: Total number of discrete feeding events
- **Feed duration**: Cumulative time spent feeding (s)
- **Meal size**: Volume consumed per individual feeding event (nL)
- **Meal duration**: Length of individual feeding events (s)
- **Feed speed**: Rate of liquid consumption during feeding events (nL/s)

Locomotor behavior was characterized using the following metrics :

- **Latency**: Time elapsed before first feeding event (s)
- **Pre-feed speed**: Mean velocity during 120 s preceding feeding events (mm/s)
- **During-feed speed**: Mean velocity during feeding events (mm/s)
- **Post-feed speed**: Mean velocity during 120 s following feeding events (mm/s)
- **Mean speed**: Average velocity across entire assay period (mm/s)
- **Mean height/elevation**: Average Y-position during assay period (mm)
- **Food port occupancy**: Percentage of time spent in food-containing feed ports
- **Control port occupancy**: Percentage of time spent in feed ports without food
- **Falls**: Number of rapid vertical displacement events

Additional behavioral parameters were calculated:

- **During-feed speed ratio**: Ratio of during-feed speed to pre-feed speed
- **Peri-feed speed ratio**: Ratio of post-feed speed to pre-feed speed

### Data analyses: estimation of effect sizes

Data analysis and interpretation were based on effect sizes (Ho et al. 2019). Summary measures for each group were plotted as a vertical line with a gap, with the ends of the line corresponding to the standard deviation and gaps corresponding to the means. Effect sizes were reported for each driver line as the mean difference between controls and test animals for all behavioral metrics. Two controls (with driver or responder genes) were grouped together, and the mean of the two controls was used to calculate the difference between the control and test flies. The mean differences and their 95 % confidence intervals (95CI) are depicted as black dots with error bars. Where possible, each error bar is accompanied by a filled curve displaying the distribution of mean differences, as calculated by bootstrap resampling. Bootstrapped distributions are robust for non-normal data. Textually, they are presented as “mean [95CI lower bound, upper bound]”. Interpretation focused on three integrated elements: (1) whether the 95 % CI overlapped zero—if not, this indicated credible evidence for an effect beyond background variation; (2) the magnitude of the point estimate and width of the CI, reflecting both biological importance and estimation precision; and (3) the shape of the distribution curve, where narrow, tall curves displaced from zero indicated robust, precise effects, while broad, shallow curves indicated greater uncertainty. Although not used for interpretation, P values were calculated by the permutation test in SciPy and are presented pro forma only; no significance tests were conducted.

### Data analysis: estimation with two variables

Our feeding experiments investigate the effects of two interacting independent variables: genotype status and light conditions on dependent variables: the relevant feeding metrics. The **delta-delta** effect size enables us to generate the net effect of these two variables (Lu et al. 2026).

The mean difference between the genotype status is:

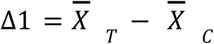

*T* is Test and *C* is Control.

The mean difference between the light on and light off conditions is:

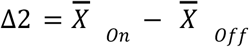

*On* is Light on and *Off* is Light off.

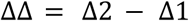

where *X̅* is the sample mean, Δ is the mean difference.

We have introduced delta g (Δg), to standardize delta-delta effects (https://acclab.github.io/DABEST-python/). Standardized mean-difference statistics, such as Hedges’ g, quantify effect sizes relative to sample variance and correct for small sample bias. This metric enables comparisons across measurements with different dimensions.

According to conventional interpretations, Δg = 0.2 represents a small effect, Δg = 0.5 a medium effect, and Δg = 0.8 a large effect.

The standard deviation of the delta-delta value is calculated from a pooled variance of the 4 samples:

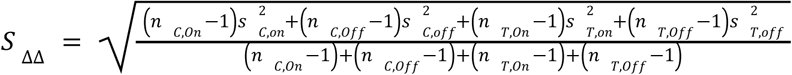

where *s* is the standard deviation and *n* is the sample size.

A delta g value is then calculated as delta-delta value divided by pooled standard deviation:

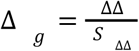

### Data analysis: meta analysis

DABEST mini-meta analysis (Lu et al. 2026) was used to calculate a weighted ΔΔ for the summary effect from a series of experiments in **Figure 3F**.

### Data analysis: linear regression

For linear regression, the Pearson correlation coefficient (r), coefficient of determination (*R^2^*), and slope were calculated with the SciPy library of Python. The pairplot scatterplots were generated with the Seaborn data visualization library (Waskom 2021).

### Data analysis: location heatmaps

The z score of time spent at different locations in the fly chamber was calculated for heat maps, and was calculated as:

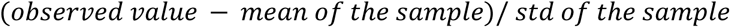

The analysis codes used for this study are available from Github repositories: https://github.com/sangyu/esploco, https://github.com/ACCLAB/espresso, https://github.com/ACCLAB/DABEST-python.

### Data analysis: time-series data

Time series data were analyzed and visualized as group means with 95% confidence intervals. Raw data were binned into 60-second intervals to reduce noise and facilitate visualization of temporal trends. For each time bin, the mean value was calculated for each individual fly, and these individual means were then averaged across flies within each experimental group to obtain the group mean at each time point. The shaded ribbons represent these 95% confidence intervals, providing a measure of variability around the group mean throughout the assay duration.

### Connectomics visualization

Synaptic connectivity between VPM and MBON11 neurons and PPL101 and MBON11 neurons, was visualized using the neuPrint skeleton viewer in the neuPrint platform, using the hemibrain v1.2.1 connectome data set (Plaza et al. 2022).

### Immunohistochemistry, GRASP, and confocal imaging

Immunohistochemistry was performed as previously described (Mohammad et al. 2017). In brief, 5–10 days male fly brains were dissected in cold phosphate buffered saline (PBS), before fixation in 4 % paraformaldehyde (Electron Microscopy Sciences) in PBT (PBS supplemented with 0.3 % Triton X-100) for 25 min. Next, the samples were washed three times with PBT (10 min each) and blocked with 5 % normal goat serum in PBT for 1 h.

Primary antibodies were applied to the samples and left to incubate for 48 h at 4°C. The samples were washed with PBT three times and incubated with secondary antibodies overnight at 4°C. Then, the brains were transferred to a slide, mounted in Vectashield mounting medium (Vectashield Vibrance® Antifade Mounting Medium; H-1700) and covered with a coverslip after washing three times in PBT. The following primary and secondary antibodies were used in the procedure: mouse anti-DLG1 (4F3 anti-DISCS LARGE 1, Developmental Studies Hybridoma Bank, 1:200 dilution) (Parnas et al. 2001), chicken anti-GFP antibody (ab13970, 1:1000), Alexa Fluor 568 goat anti-mouse (Invitrogen 1:500), Alexa Fluor 488 goat anti-chicken (Invitrogen 1:1000). To measure native GRASP-induced GFP signals (Gordon and Scott 2009), adult male flies were collected 5–10 days after eclosion and their brains were dissected in cold PBS and fixed in 4 % PFA in PBS for 20 min at 23 °C and washed three times in PBT (10 min each). The GRASP brains were mounted in Vectashield medium (Vectashield Vibrance® Antifade Mounting Medium; H-1700) without antibody staining.

Confocal images of the brain samples were captured under an LSM710 Carl Zeiss confocal microscope. The maximum-intensity projections of the confocal images were generated with the FIJI distribution of ImageJ (Schindelin et al. 2012) and the colors were inverted using the EZreverse web application (Song and Goedhart 2024).

## Acknowledgments

We thank Dr. Sarah J Certel for sharing the *Tdc2-lexA:VP16* line. We thank Dr Ravinuthula Sruthi Jagannathan (Duke-NUS Medical School) and Dr Jessica Tamanini (Insight Editing London) for their critical reviews of the manuscript before submission. We thank the Duke-NUS Medical School core imaging facility. We thank Gerry Rubin of Janelia Research Campus for sharing fly lines. The 4F3 anti-Discs large monoclonal antibody was obtained from the Developmental Studies Hybridoma Bank, created by the NICHD of the NIH and maintained at The University of Iowa, Department of Biology, Iowa City, IA 52242. Stocks obtained from the Bloomington Drosophila Stock Center (NIH P40OD018537) were used in this study. Transgenic fly stocks and/or plasmids were obtained from the Vienna Drosophila Resource Center (VDRC) at Vienna BioCenter Core Facilities (VBCF), member of the Vienna BioCenter (VBC), Austria.

## Author Contributions

**Conceptualization**: XY, SX, ACC; **Experiment design**: XY, SX, ACC; **Methodology**: XY, SX; **Software**: SX, JH (Python), JCS (CRITTA, LabView); **Data Analysis:** XY, SX (Python); **Investigation**: XY (feeding assays, genetics, brain dissection, immunohistochemistry, and microscopy); SX (feeding assays) **Writing – Original Draft:** XY; **Writing – Revision:** XY, ACC, SX; **Visualization:** XY, SX, ACC, JH; **Supervision:** ACC; **Project Administration:** ACC; **Funding Acquisition:** ACC.

